# Metabolic engineering of *Corynebacterium glutamicum* for the production of anthranilate from glucose and xylose

**DOI:** 10.1101/2023.06.24.546385

**Authors:** Mario Mutz, Vincent Brüning, Christian Brüsseler, Moritz-Fabian Müller, Stephan Noack, Jan Marienhagen

## Abstract

Anthranilate and its derivative aniline are important basic chemicals for the synthesis of polyurethanes as well as various dyes and food additives. Today, aniline is mainly chemically produced from petroleum-derived benzene, but it could be also obtained more sustainably by decarboxylation of the microbially produced shikimate pathway intermediate anthranilate. In this study, *Corynebacterium glutamicum* was engineered for the microbial production of anthranilate from a carbon source mixture of glucose and xylose. First, a feedback-resistant 3-deoxy- arabinoheptulosonate-7-phosphate synthase from *E. coli*, catalyzing the first step of the shikimate pathway, was functionally introduced into *C. glutamicum* to enable anthranilate production. Modulation of the translation efficiency of the genes for the shikimate kinase (*aroK*) and the anthranilate phosphoribosyltransferase (*trpD*) improved product formation. Deletion of two genes, one for a putative phosphatase (*nagD*) and one for a quinate/shikimate dehydrogenase (*qsuD*), abolished by-product formation of glycerol and quinate. However, the introduction of an engineered anthranilate synthase (TrpEG) unresponsive to feedback inhibition by tryptophan had the most pronounced effect on anthranilate production. Component I of this enzyme (TrpE) was engineered using a biosensor-based *in vivo* screening strategy for identifying variants with increased feedback-resistance in a semi-rational library of TrpE muteins. The final strain accumulated up to 5.9 g/L (43 mM) anthranilate in defined CGXII medium from a mixture of glucose and xylose in bioreactor cultivations. We believe that the constructed *C. glutamicum* variants are not only limited to anthranilate production, but could also be suitable for the synthesis of other biotechnologically interesting shikimate pathway intermediates, or any other aromatic compound derived thereof.

## 1 Introduction

Aromatic compounds have a wide range of applications in the chemical, pharmaceutical, or food industries where they serve as polymer building blocks, dyes, food additives, or antibiotics (Balderas-Hernández *et al*., 2009; Chung, 2016; Noreen *et al*., 2016). However, the chemical production of these compounds at an industrial scale is typically based on benzene, toluene, or xylene (BTX) derived from crude oil (Haveren *et al*., 2007). One of the key aromatic platform chemicals is aniline with a global market volume of 9.4 million tons in 2022, which is expected to increase to more than 12 million tons by 2028 with an annual growth rate of 5.2 % (IMARC Group 2023: “Aniline Market: Global Industry Trends, Share, Size, Growth, Opportunity and Forecast 2023-2028”). Three quarters of the aniline produced globally is used for the synthesis of methylene diphenyl diisocyanate (MDI), a monomer building block for polyurethane (PU), which in turn is a polymer utilized in the manufacturing of foam, elastomers, paints, adhesives or artificial leather (Driessen *et al*., 2017; Nakajima-Kambe *et al*., 1999). Aniline is mainly produced chemically from benzene, which is initially converted to nitrobenzene with nitric acid. (Bolden *et al*., 2015). Subsequently, the nitro functionality is reduced to the primary amine in the gas phase at 270°C and 1.25 bar employing a copper catalyst. This process releases 25 Mt CO_2_ and consumes 0.3 EJ energy annually (Winter *et al*., 2021).

However, aniline could be also produced in a more sustainable manner using the metabolic capabilities of microorganisms. This biotechnological process involves the microbial production of anthranilate (ANT), which can be subsequently converted to aniline by chemical decarboxylation. Besides aniline, ANT is also the precursor of other aromatic compounds of biotechnological interest such as methyl anthranilate (MANT) conveying the typical scent and taste of the Concord grape (*Vitis labrusca*) (Chambers *et al*., 2013; Fuleki, 1972; Wang & Luca, 2005). Another derivative is menthyl anthranilate, which serves as a UV-A quencher in sunscreens (Beeby & Jones, 2000). ANT is an intermediate of the shikimate (SA) pathway, which is responsible for the synthesis of the three aromatic amino acids (TYR, PHE, TRP), as well as secondary metabolites in plants, fungi, and many microorganisms (Lee & Wendisch, 2017) (Fig. 1).

**Fig. 1:**
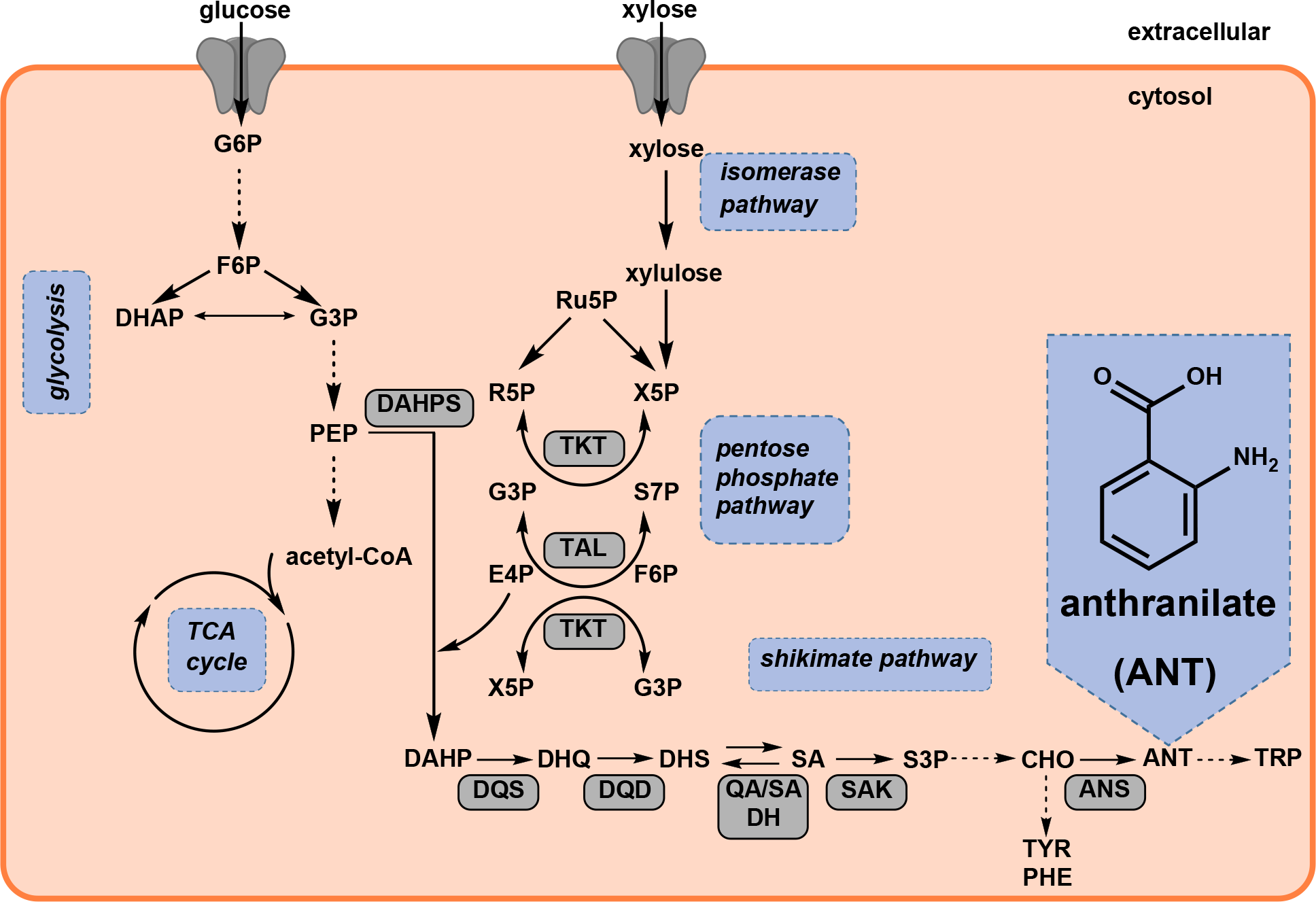
Biosynthesis of anthranilate from glucose and xylose in *C. glutamicum*. acetyl-CoA, acetyl-coenzyme A; ANS, anthranilate synthase; ANT, anthranilate; CHO, chorismate; DAHPS, 3-deoxy- arabinoheptulosonate-7-phosphate (DAHP) synthase; DHAP, dihydroxyacetone phosphate; DHQ, 3- dehydroquinate; DHS, 3-dehydroshikimate; DQD, 3-dehydroquinate dehydratase; DQS, 3-dehydroquinate synthase; E4P, erythrose-4-phosphate; F6P, fructose-1,6-bisphosphate; G3P, glycerinaldehyde-3- phosphate; G6P, glucose-6-phosphate; PEP, phosphoenolpyruvate; PHE, phenylalanine; QA/SA DH, quinate/shikimate dehydrogenase; R5P, ribose-5-phosphate; Ru5P, ribulose-5-phosphate; SA, shikimate; SAK, shikimate kinase; S3P, shikimate-3-phosphate; S7P, sedoheptulose-7-phosphate; TAL, transaldolase; TKT, transketolase; TCA cycle, tricarboxylic acid cycle; TRP, tryptophan; TYR, tyrosine; X5P, xylulose-5-phosphate.

The first step in the synthesis of aromatic compounds via the SA pathway is the condensation of erythrose-4-phosphate (E4P), a metabolite of the non-oxidative part of the pentose phosphate pathway (PPP), and phosphoenolpyruvate (PEP), an intermediate of glycolysis, yielding 3-deoxy-arabinoheptulosonate-7-phosphate (DAHP) (Herrmann & Weaver, 1999). The reaction is catalyzed by DAHP synthase, which is allosterically regulated (Liao *et al*., 2001; Shumilin *et al*., 1996). After seven enzymatic conversions, the SA pathway bifurcates at the stage of chorismate, eventually leading to PHE, TYR and TRP. The first committed step towards TRP biosynthesis is catalyzed by the anthranilate synthase (ANS) converting chorismate to ANT and pyruvate, using GLN as an amino donor (yielding GLU) (Romero *et al*., 1995).

The microbial production of ANT was established in a few bacterial species. A *Pseudomonas putida* KT2440 strain accumulated 1.54 g/L (11.23 mM) ANT in a defined medium with glucose as a carbon source using a fed-batch process (Kuepper *et al*., 2015). In addition, a plasmid-free strain of *P. putida* was constructed, which produced 3.8 mM ANT in shake flasks from glucose without the supplementation of additives (Fernández-Cabezón *et al*., 2022). Higher ANT titers were only achieved by using expensive complex media. A *Bacillus subtilis* strain produced 3.5 g/L (25 mM) ANT using yeast extract and glucose as substrates within 60 h, whereas an engineered *Escherichia coli* variant accumulated 14 g/L (102 mM) ANT after 34 h in M9 minimal medium supplemented with 10 g/L of yeast extract and 30 g/L glucose followed by two pulses containing the same carbon source composition (Balderas-Hernández *et al*., 2009; Cooper *et al*., 1995). Moreover, a titer of 567.9 mg/L (4.1 mM) ANT was achieved with *Saccharomyces cerevisiae* using a complex SCD medium, which is the highest ANT titer reached using a eukaryotic host (Kuivanen *et al*., 2021). Furthermore, MANT was produced from ANT with *C. glutamicum* using fed-batch cultures and a two-phase cultivation mode with a maximum titer of 5.74 g/L (38 mM) MANT (Luo *et al*., 2019). In the context of this study, the first ANT-production with *C. glutamicum* was described. The best variant accumulated up to 26.4 g/L (192.5 mM) ANT. However, to achieve such high product titers. the ANT-producing *C. glutamicum* strain was made auxotrophic for TRP and thus required TRP-supplementation every 24 h.

ANT, similar to most other aromatics, is known to have antimicrobial properties, potentially limiting the microbial production of this compound (Li *et al*., 2017). *C. glutamicum* is used for the industrial production of proteinogenic amino acids such as GLU and LYS at a scale of more than five million tons per year (Eggeling & Bott, 2005). This gram-positive soil bacterium is free of endotoxins and products derived from *C. glutamicum* are therefore generally recognized as safe (Zha *et al*., 2018). In addition, several *C. glutamicum* variants engineered for aromatic compounds such as hydroxybenzoic acids, phenylpropanoids, or plant polyphenols are available demonstrating that *C. glutamicum* is a robust host system capable of enduring increased concentrations of cytotoxic aromatics (Kallscheuer *et al*., 2016; Kallscheuer & Marienhagen, 2018; Milke *et al*., 2019; Son *et al*., 2021).

In addition to a wide product range, *C. glutamicum* can metabolize various cheap carbon sources. Frequently, glucose and fructose are used as carbon and energy source for applied purposes (Blombach & Seibold, 2010). However, as the utilization of these sugars in biotechnological processes competes with food production, it is highly desirable to resort to other substrates, preferably from waste streams, such as cellobiose, arabinose, and xylose (Jiang & Zhang, 2016; Kawaguchi *et al*., 2006; Kotrba *et al*., 2003; Schneider *et al*., 2011). Since *C. glutamicum* cannot utilize xylose naturally, several different pathways such as the isomerase pathway or the Weimberg pathway were implemented into the metabolism of this bacterium (Kawaguchi *et al*., 2006; Meiswinkel *et al*., 2013; Radek *et al*., 2014). In this context, *C. glutamicum* was engineered to produce several compounds such as protocatechuate, succinate, or α-ketoglutarate from xylose or glucose/xylose mixtures (Brüsseler *et al*., 2019; Labib *et al*., 2021; Tenhaef *et al*., 2021). Noteworthy, xylose utilization via the isomerase pathway has the advantage of providing the SA pathway precursor E4P (Kogure *et al*., 2016).

This study aimed to establish microbial ANT production with *C. glutamicum* from different carbon sources (and mixtures) by combining rational metabolic engineering strategies with biosensor-guided semi-rational enzyme engineering.

## 2 Material and Methods

### 2.1 Bacterial strains, plasmid construction, media

Strains and plasmids utilized in this study are listed in Tab. S1. *C. glutamicum* strains were cultivated aerobically at 30°C in brain heart infusion (BHI) medium (Difco Laboratories, Detroit, USA) or in defined CGXII medium supplemented with either 4 % (w/v) glucose or a mixture of 1 % (w/v) glucose and 3 % (w/v) xylose as sole carbon and energy source (Keilhauer *et al*., 1993).

For cultivation of *C. glutamicum*, test tubes with 5 mL BHI medium supplemented with the appropriate antibiotic were inoculated with a single colony from a fresh BHI agar plate. These first precultures were grown for 8 h on a rotary shaker at 170 rpm. The entire precultures were used to inoculate the second precultures, which consisted of 50 mL defined CGXII medium supplemented with the respective carbon source in 500 mL baffled flasks. These second precultures were cultivated overnight on a rotary shaker at 130 rpm. Bacterial growth was tracked by measuring the optical cell density at 600 nm (OD_600_). CGXII main cultures with the indicated carbon source were inoculated to an OD_600_ of 1 from these second precultures. Strains utilized in this study are equipped with the chromosomally integrated gene *1* encoding the T7 RNA polymerase enabling gene expression from the IPTG-inducible T7 promotor (Kortmann *et al*., 2015). Thus, for ANT production, heterologous and homologous gene expression was induced 1 h after inoculation with 20 µM IPTG. At defined time points, 1 mL of the culture broth was sampled, and centrifuged at 13,300 rpm for 20 min and the culture supernatant was stored at - 20°C until HPLC analysis.

To determine the cytotoxic effect of ANT on bacterial growth, *C. glutamicum* was cultivated at 30°C, 900 rpm, and a humidity of 85 % in 48-well Flowerplates containing 800 µL defined CGXII medium with 0.5 % (w/v) glucose inoculated to an OD_600_ of 1 by using the BioLector microbioreactor for 24 h (Beckman Coulter Life Sciences, Aachen, Germany). Increasing concentrations of ANT (final concentration of 0, 0.5, 1, 2.5, 5, 7.5, 10, and 12 g/L) were added to the medium. A 200 g/L ANT stock solution, dissolved in H_2_O was prepared and the pH was adjusted to 7 by using a 1 M NaOH solution. As the stock solution could not be concentrated higher due to the low solubility of ANT in water (5.7 g/L (41.7 mM) at 25 °C), the maximum ANT concentration in defined CGXII medium was 12 g/L. To monitor bacterial growth, the backscattered light intensity (620 nm, gain 10) was followed.

*E. coli* DH5α was utilized for molecular cloning and plasmid propagation, while *E. coli* TOP10 was used for the construction of strain libraries. Cultivation of *E. coli* strains was routinely performed in Lysogeny Broth (LB) medium (10 g/L tryptone, 10 g/L NaCl, 5 g/L yeast extract) or yeast nitrogen base (YNB) medium at 37 °C (Bertani, 1951). For the preparation of 1 L YNB medium, 100 mL 10x concentrated YNB supplemented with 5.1 % (v/v) glycerol was added to 900 mL YNB base (6 g/L K_2_HPO_4_, 3 g/L KH_2_PO_4_, 10 g/L 3-(N-morpholino)propanesulfonic acid (MOPS) pH 7). For the LEU auxotroph *E. coli* DH10B-derived strains, 2 mM LEU was supplemented from a 20 mM LEU stock solution in the YNB base (Durfee *et al*., 2008). Where necessary, kanamycin (*E. coli*: 50 µg/mL in LB medium and 25 µg/mL in YNB medium; *C. glutamicum*: 25 µg/mL for cultivations, 15 µg/mL for colony selection after transformation) and/or spectinomycin (100 µg/mL for both, *E. coli* and *C. glutamicum*) was added to the medium.

### 2.2 Bioreactor cultivations

First precultures were inoculated from cells grown on a fresh BHI agar plate in 10 mL BHI medium using 100 mL baffled flasks. These precultures were incubated at 30°C and 250 rpm for 6 h. Subsequently, these precultures were split and 5 mL of each preculture was used for inoculation of two second precultures, which consisted of 100 mL defined CGXII medium supplemented with either 1 % glucose and 3 % xylose (batch mode) or 4 % glucose (fed-batch mode) in 1 L baffled flasks, which were incubated at 30 °C and 250 rpm for 24 h. For inoculation of the main culture, the second precultures were centrifuged for 10 min at 4,000 rpm and 4 °C. The supernatants were discarded and the cells were washed in 20 mL 0.9 % NaCl. The cells of the two second precultures were merged, the washing step was repeated and the cells were resuspended in 5 mL 0.9 % NaCl. Finally, OD_600_ was measured and bioreactors were inoculated to an initial OD_600_ of 1.

Batch and pulsed-fed-batch bioreactor fermentations were performed in duplicates in DASGIP bioreactor systems (Eppendorf, Hamburg, Germany) with a total volume of 2 L and an initial working volume of 1 L with four simultaneously operated reactors. Cells were cultivated in defined CGXII medium without urea and MOPS supplemented with 1 % glucose and 3 % xylose during the batch cultivations. For the fed-batch cultivations, 4 % glucose for initial biomass formation was used as a carbon source. After 14.5 h, glucose was depleted and a constant glucose feed started with a rate of 0.5 g/h. Simultaneously, the first 10 g xylose pulse was added from a 50 % xylose stock solution. Subsequently, xylose was pulsed after 18.75 h (10 g), 21.75 h (10 g), 37.42 h (10 g), 41.42 h(10 g), 45.42 h (20 g), 62.83 h (10 g), 66.25 h (10 g) and 69.75 h (20 g). After inoculation, 2 mL sterile antifoam was added and plasmid-based gene expression was induced with 20 µM IPTG after 1 h. The temperature was set to 30 °C and the pH was maintained at 7 by both-sided regulation with 5 M H_2_PO_4_ and 25 % NH_4_OH. Dissolved oxygen (dO_2_) was fixed to a minimum of 30 % using a cascade. The cascade first increased agitation from 400 to 1,200 rpm, followed by the airflow, which was increased from 6 to 40 sL/h if necessary. The cultivations were terminated after the complete depletion of the carbon sources. Samples were taken at defined time points, centrifuged at 13,300 rpm for 20 min and the supernatant was stored at -20 °C until ANT, glucose, and xylose quantification via HPLC. Data from bioreactor experiments in batch mode were modeled using Monod kinetics to describe growth of biomass, consumption of glucose and xylose consumption, and production of anthranilate. Specific dynamics of xylose utilization were modelled by considering additional inhibition by glucose. For model implementation, validation and analysis we used the open-source, python-based modeling tool pyFOOMB (Hemmerich *et al*., 2021).

### 2.3 Plasmid and strain construction

Standard procedures such as molecular cloning, PCR, and DNA restriction and ligation were performed as described elsewhere (Sambrook *et al*., 2001). Genes and chromosomal fragments required for plasmid construction were amplified by PCR using genomic *C. glutamicum* DNA or whole cells as a template. Oligonucleotides are listed in Tab. S2. PCR fragments were cloned into plasmid vectors by Gibson assembly (Gibson *et al*., 2009). Gene deletions and the introduction of point mutations or whole genes in the genome of *C. glutamicum* were achieved by employing a two-step homologous recombination method described previously (Niebisch & Bott, 2001; Schäfer *et al*., 1994). Colony PCR, restriction analysis, and DNA sequencing performed at Eurofins MWG Operon (Ebersberg, Germany) were used to verify plasmid constructions and the identity of recombinant *C. glutamicum* strains. Transformation of *C. glutamicum* with constructed plasmids was performed by electroporation (Kirchner & Tauch, 2003).

For site-saturation mutagenesis (SSM), plasmid pJC1-*trpE* (7749 bp) was amplified by PCR using oligonucleotides with degenerated (NNK) codons (NEB Q5 Site-Directed-Mutagenesis Kit, New England Biolabs, GmbH, Frankfurt am Main, Germany). PCRs were performed using the Q5 polymerase (Q5 Hot Start High-Fidelity, 25 ng template, 10 µM of each primer, 30 cycles, 30 s initial denaturation, 10 s denaturation, 30 s annealing, 433 s extension, 120 s final extension). After PCR amplification, the parental plasmid was removed and the plasmids were constructed by a kinase-ligase-*Dpn*I (KLD) reaction according to the instructions of the manufacturer. The resulting plasmid libraries were used for heat shock transformation of chemically competent One Shot TOP10 *E. coli* cells. All resulting transformants were detached from LB agar plates and used for inoculation of 50 mL LB medium in 500 mL baffled shake flasks, which were cultivated on a rotary shaker at 37° for 6 h before plasmid isolation using a Midi Kit (Qiagen, Hilden, Germany). The obtained plasmid libraries were subsequently used for electroporation of *C. glutamicum* ANT5 Δ*trpE*.

### 2.4 Biosensor-based screening

From each of the four generated ANS libraries, 150 variants were screened for improved ANT production. Initially, 800 µL BHI + 1 % (w/v) glucose (BHIG) glycerol cultures of all variants were prepared in organic solvent-resistant deep-well plates. Due to the deletion of the genomic *trpE* gene, the strains were TRP auxotroph, so 0.5 mM TRP was always supplemented. By following this strategy, similar growth of all cultivated variants was ensured. The cultivations were performed in a Multitron Pro HT Incubator (Infors AG, Bottmingen, Switzerland) at 30 °C, 900 rpm, 75 % humidity, and 3 mm throw for 16 h. After the addition of 30 % (v/v) glycerol, the *C. glutamicum* strain variants were stored at -80 °C until use.

50 µL of fresh glycerol cultures were used for inoculation of BHIG precultures in 48-well Flowerplates with a total volume of 800 µL. The cultivations were performed as described above for 8 h. The second precultures (CGXII medium + 4 % (w/v) glucose + 0.5 mM TRP) were inoculated with 50 µL of the grown BHIG precultures to a total volume of 800 µL per well and incubated in the same shaker overnight. The main cultures (CGXII + 4 % glucose) were inoculated with 50 µL of the grown CGXII precultures and incubated for 72 h under the same conditions. To harvest culture supernatants containing ANT, the entire cultures were transferred into organic solvent-resistant deep-well plates, which were subsequently centrifuged (3,500 rpm, 20 min, 4°C). Cell-free culture supernatants were stored at -20°C until their biosensor-based characterization. For the correlation of specific fluorescence and ANT concentration, selected variants displaying higher, equal, or lower specific fluorescence compared to the control were also analyzed by HPLC (Tab. S3-5).

For the semi-quantitative conversion of the ANT concentration to a fluorescence signal, the strain *E. coli* DH10B Δ*hcaREFCBD* was used. Single colonies of the sensor strain *E. coli* DH10B Δ*hcaREFCBD* pSen6MSA were picked from fresh LB agar plates for inoculation of test tubes filled with 5 mL LB medium. These precultures were grown for 8 h on a rotary shaker at 170 rpm. The precultures were used to inoculate second precultures of 50 mL YNB medium, 0.51 % glycerol, and 2 mM LEU in 500 mL baffled flasks. The second precultures were cultivated overnight on a rotary shaker at 130 rpm and used for the inoculation of 50 mL YNB medium, 0.51 % glycerol, and 2 mM LEU (main culture) to an OD_600_ of 1. The inoculated main cultures supplemented with *C. glutamicum* ANT5 Δ*trpE* pJC1-*trpE*-x culture supernatant were transferred into 48-well Flowerplates to a final volume of 800 µL. The cells were cultivated in a Multitron Pro HT Incubator (Infors AG, Bottmingen, Switzerland) at 37 °C, 900 rpm, 75 % humidity, and 3 mm throw for 24 h. The supernatant of the strain equipped with wild-type *trpE* gene in biological triplicates served as control. For calibration, 0, 0.15, 0.3, 0.6, 1.25, 2.5, and 5 mM ANT were added separately from a 20 mM ANT stock solution. For blank measurements, YNB medium without cells was used. On the next day, 100 µL of each well was transferred to a 96-well Flat-bottom plate (Brand GmbH + Co. KG, Wertheim, Germany). For the biosensor-based characterization of *C. glutamicum* strain variants, fluorescence, and absorbance were measured using a plate reader (Tecan Infinite 200 PRO, Tecan Group, Maennedorf, Switzerland) and the program *YFP in cells* (λ_Ex_ = 503 nm; λ_Em_ = 540 nm, λ_Abs_ = 600 nm). The fluorescence and absorbance of the YNB medium without cells were subtracted from all fluorescence and absorbance signals. The specific fluorescence was calculated as the ratio of the fluorescence signal and absorbance at 600 nm.

### 2.5 GC-TOF-MS and HPLC analysis

The qualitative and quantitative detection of metabolites in culture supernatants was performed by high-performance-liquid-chromatography (HPLC) using a 1260 Infinity II System equipped with a 1260 Infinity II Diode Array Detector (DAD) (Agilent Technologies, Waldbronn, Germany). Samples of the culture supernatant were thawed and centrifuged for 20 min to remove cells and precipitated media components. Typically, samples taken after 8 h of cultivation were diluted (1:5) with ddH_2_O in order to ensure a metabolite concentration within the linear range of the authentic standard. Authentic standards of ANT, SA, quinate (QA), glycerol, glucose, and xylose were purchased from Sigma-Aldrich (Taufkirchen, Germany). For the isocratic separation of ANT, an Agilent InfinityLab Poroshell 120 2.7 µM EC-C_18_ column (3.0 x 150 mm; Agilent Technologies, Waldbronn, Germany) at 40°C with an InfinityLab Poroshell 2.7 µM EC-C_18_ pre-column (3 x 5 mm; Agilent Technologies, Waldbronn, Germany) was used. For elution from the column, 0.1 % (v/v) acetic acid (solvent A, 80 %) and acetonitrile supplemented with 0.1 % acetic acid (solvent B, 20 %) were applied as mobile phase at a flow rate of 0.35 mL/min. The concentration of ANT was determined by measuring the absorption at 220 nm.

For the isocratic separation of organic acids and sugars, a Rezex ROA-Organic Acid H^+^ column 8 µm (300 x 7.8 mm; Phenomenex, Torrance, California, USA) in combination with a Security Guard HPLC Guard Cartridge system Carbo-H 4 x (4 x 3 mm; Phenomenex, Torrance, California, USA) as pre-column was used at 80 °C. For elution from the column, 5 mM H_2_SO_4_ was used as the mobile phase at a flow rate of 0.3 mL/min. The concentration of QA and SA was determined by measuring the absorption at 220 nm, while glycerol, glucose, and xylose were detected by the refractive index detector (RID). Area values of integrated signals were linear up to metabolite concentration of 5 mM (ANT), 10 mM (SA, QA, glycerol), or 100 mM (glucose, xylose).

Gas chromatography-time-of-flight (GC-TOF) mass spectrometry (MS) was performed for metabolite identification in culture supernatants using an Agilent 8890N double SSL gas chromatograph (Agilent, Waldbronn, Germany) equipped with a L-PAL3-S15 liquid autosampler coupled to a LECO GCxGC HRT+ 4D high resolution TOF MS (LECO, Mönchengladbach, Germany). Sample preparation, two-step derivatization of the samples, and MS data acquisition were performed as described previously (Paczia *et al*., 2012). Peak identification of known and unknown metabolites was performed as described before (de Witt *et al*., 2023).

## 3 Results

### 3.1 Cytotoxicity of anthranilate for *C. glutamicum*

Previously, the platform strain *C. glutamicum* DelAro^5^ PO6-*iolT1* was constructed, which is devoid of a large portion of the extensive catabolic network for aromatic compounds (Kallscheuer *et al*., 2016). This strain served as starting strain for rational metabolic engineering to establish microbial ANT synthesis with *C. glutamicum*. However, prior to any engineering, the robustness of *C. glutamicum* to ANT was investigated. For these experiments, *C. glutamicum* DelAro^5^ P_O6_- *iolT1* was cultivated in the presence of 0-12 g/L ANT in microtiter plates (MTPs) (Fig. 2A). These experiments showed that the growth of *C. glutamicum* DelAro^5^ P_O6_-*iolT1* was negatively affected with increasing ANT concentrations starting from the lowest concentration of 0.5 g/L (3.65 mM) ANT. ANT supplementation resulted in a prolonged lag phase and a reduced growth rate (Fig. 2B). At the highest concentration of 12 g/L (87.5 mM) ANT, the strain was still able to grow with a maximal growth rate of 0.1 h^-1^, which is 50 % of the maximal growth rate determined for the control (µ_max_ = 0.2 h^-1^) indicating that *C. glutamicum* DelAro^5^ P_O6_-*iolT1* is capable of tolerating high ANT concentrations.

**Fig. 2:**
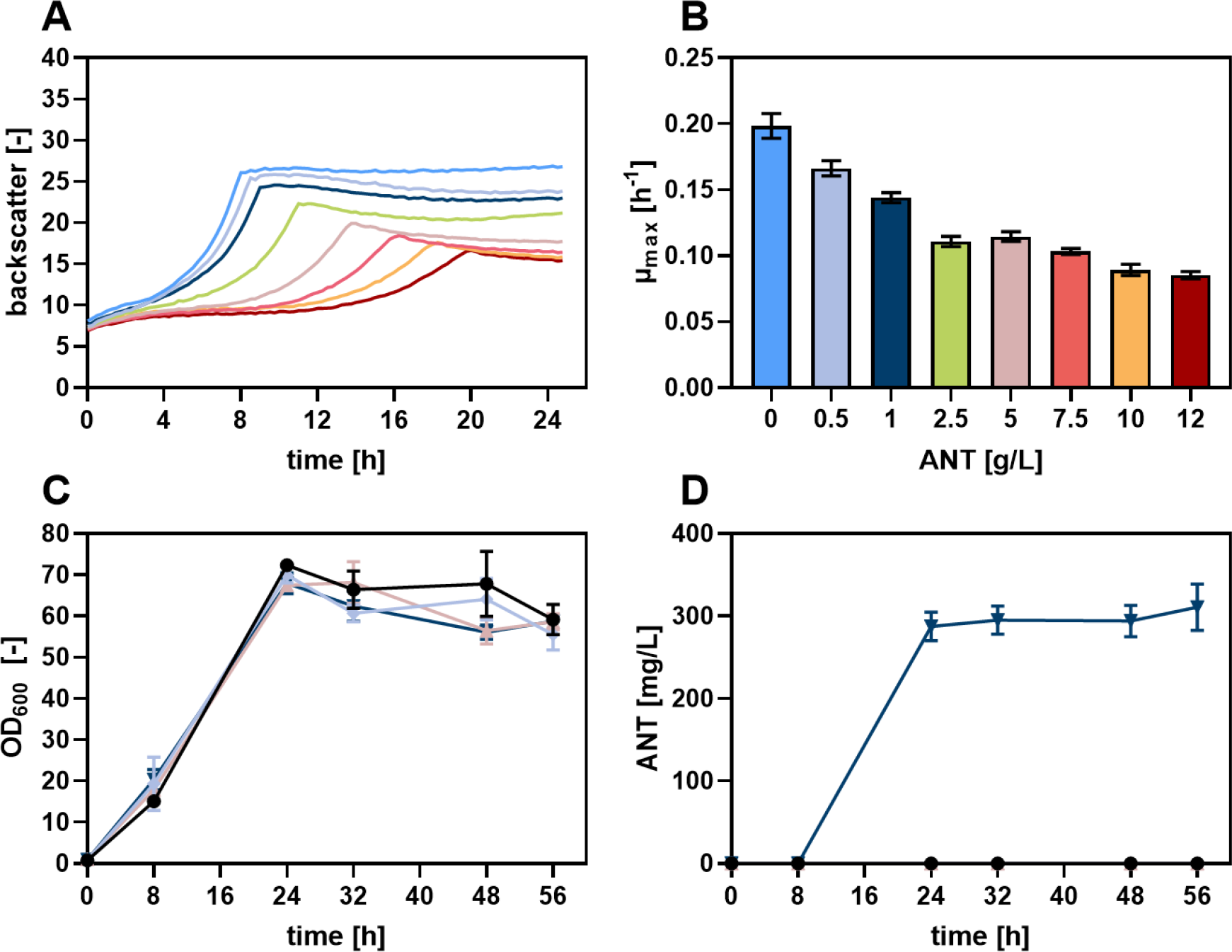
**Investigation of ANT toxicity and ANT synthesis with *C. glutamicum***. (**A**) Cytotoxic effects of ANT on the growth of *C. glutamicum* DelAro^5^ P_O6_-*iolT1*. For a better visibility, only the respective mean values were displayed for the growth curves. (**B**) Growth rates for each ANT concentration tested. The depicted data represent the mean values and standard deviation of six individual experiments (n = 6). (**C**) Growth (OD_600_) of *C. glutamicum* DelAro^5^ P_O6_-*iolT1* (circles), *C. glutamicum* ANT1 (diamonds), *C. glutamicum* ANT1 pMKEx2 (triangles), and *C. glutamicum* ANT1 pMKEx2-*aroF*\**_EcCg_*-*tkt_Cg_* (reversed triangles). (**D**) Determination of the ANT titer in the culture supernatants by HPLC. The depicted data represent mean values and standard deviation of biological triplicates.

### 3.2 Establishing microbial anthranilate production from glucose

Heterologous expression of a gene encoding a feedback-resistant DAHP synthase is an established strategy for the microbial production of SA-derived compounds (Balderas-Hernández *et al*., 2009). In this study, the codon-optimized gene *aroF*\**_EcCg_* encoding a DAHP synthase from *E. coli* was selected, which was previously engineered to be feedback-resistant to TYR (Zhang *et al*., 2014). Furthermore, increased transketolase activity was described to be associated with aromatic amino acid synthesis due to the increased carbon flux through the SA pathway, providing more E4P (Ikeda, 2006). For this reason, *tkt* encoding the transketolase from *C. glutamicum* was also selected for plasmid-based homologous overexpression. With the aim to increase the translation efficiency of *tkt*, the native TTG-start codon was replaced by ATG. Both genes, *aroF*\**_EcCg_* and *tkt* were cloned as a bicistronic operon on the pMKEx2-plasmid yielding pMKEx2- *aroF*\**_EcCg_*-*tkt_Cg_*, which allows for IPTG-inducible expression under the control of the strong T7 promotor (Kortmann *et al*., 2015). This plasmid was transformed into *C. glutamicum* DelAro^5^ P_O6_- *iolT1* GTG-*trpD* (ANT1) in which the ATG-start codon of the genome-encoded *trpD*-gene was replaced by a GTG to reduce the initial translation efficiency of this gene. The *trpD*-gene product, essential for growth, is the anthranilate phosphoribosyltransferase, which converts ANT to *N*-(5- phosphoribosyl)-anthranilate and is thus withdrawing ANT for TRP synthesis (O’Gara & Dunican, 1995). The resulting strain *C. glutamicum* ANT1 pMKEx2-*aroF*\**_EcCg_*-*tkt_Cg_* and suitable control strains were cultivated in shake flasks with glucose as the sole carbon and energy source (Fig. 2C, D). Samples of the culture supernatants were analyzed for ANT production by HPLC. Neither *C. glutamicum* ANT1 nor the strain carrying the empty vector produced detectable amounts of ANT (Fig. 2D). The strain capable of plasmid-based *aroF*\**_EcCg_* - and *tkt_Cg_*-expression accumulated 287.6 ± 14.1 mg/L (2.1 mM) ANT within 24 h (Fig. 2D).

However, in addition to ANT, the accumulation of an additional compound was detected during HPLC analysis. The shorter retention time did hint towards the previously described formation of glycosyl-anthranilate in the presence of catalytically active cations such as ammonium, which is a component of the defined CGXII medium used (Kuepper *et al*., 2020). Hence, glycosyl-anthranilate or its Amadori product fructosyl-anthranilate might be formed, the latter being characterized by the same absorption spectrum in the UV range as ANT (Fig. S1A, B). Since no analytical standard of this compound was available, ANT was dissolved in CGXII medium with and without glucose and incubated at 30 °C for 72 h. HPLC analysis showed indeed that the putative glycosyl-anthranilate peak only emerged in the presence of glucose, whereas the area of the ANT peak decreased simultaneously.

Samples of the culture supernatants were also examined by GC-TOF MS to detect the possible formation of other by-products. (Fig. S2). These analyses showed that glycerol accumulated as a by-product. Since glycerol, but also QA, are known by-products typically accumulating during the microbial production of SA pathway-derived products, genes responsible for the formation of these compounds were deleted (Kogure *et al*., 2016). The gene *nagD* encodes the putative phosphatase HdpA, involved in the formation of dihydroxyacetone, the precursor of glycerol (Jojima *et al*., 2012). The gene *qsuD* encodes the QA/SA dehydrogenase QsuD, which catalyzes the first step of QA/SA catabolism (Fig. 1). By deletion of *qsuD*, the conversion of 3- dehydroquinate (DHQ) to QA and also the undesired back reaction of SA to 3-dehydroshikimate (DHS) could be prevented (Teramoto *et al*., 2009). Hence, respective deletion strains of *C. glutamicum* ANT1 were constructed, which differ from the parent strain by in-frame deletion of *nagD*, *qsuD,* or both genes. The effect on ANT production was investigated by the cultivation of all constructed deletion strains in shake flasks (Fig. S3). Neither the deletion of *nagD, qsuD* nor both genes in combination had a negative effect on growth and maximum cell density was reached after 13 h for all strains, whereupon product formation set in. This indicated a growth-decoupled formation of ANT from glucose. The product titer of the single deletion strains was not increased compared to the control. However, the double deletion strain *C. glutamicum* ANT1 Δ*nagD* Δ*qsuD* (ANT2) pMKEx2-*aroF*\**_EcCg_*-*tkt_Cg_* achieved a higher average ANT titer determined to be 294 ± 11.9 mg/L (2.2 mM) ANT (Fig. S3B).

### 3.3. Anthranilate production glucose/xylose mixtures

In order to enable ANT production also from xylose, the xylose isomerase pathway was introduced into *C. glutamicum* ANT2 pMKEx2-*aroF*\**_EcCg_*-*tkt_Cg_* via the plasmid pEKEx3-*xylA_Xc_*- *xylB_Cg_* (Meiswinkel *et al*., 2013). The *xylA_Xc_*-encoded xylose isomerase from *Xanthomonas campestris* and the *xylB_Cg_*-encoded xylulokinase from *C. glutamicum* catalyze the conversion of xylose to xylulose-5-phosphate, which is a readily metabolizable intermediate of the non-oxidative part of the PPP. This in turn could increase the availability of the DAHP synthase substrate E4P. The ability of the strains to utilize xylose as a carbon source was investigated and the effect of xylose utilization on ANT production was tested by the cultivation of *C. glutamicum* ANT2 harboring both expression plasmids in shake flasks with a mixture of glucose and xylose (Fig. S4). By using a mixed carbon substrate of 1 % glucose and 3 % xylose, the cultures reached optical densities of 50 within 20 h (Fig. S4A), while higher optical densities of 70 were reached with 4 % glucose as the sole carbon source within 13 h. However, from the glucose/xylose mixture, ANT could be produced at a g/L scale with an average titer of 2 g/L (14.6 mM) ANT (Fig. S4B). Interestingly, ANT accumulation started before the cells entered the stationary growth phase in these cultivations, whereas product formation with glucose as the sole carbon source occurred in a growth-decoupled manner. Subsequently, the effects of different glucose/xylose ratios were investigated (Fig. S5). These experiments showed that ANT formation increased with increasing concentrations of xylose and decreasing glucose concentrations. Growth and product formation from xylose as the sole carbon and energy source was possible, but did not prove to be beneficial since the strains were characterized by a prolonged lag phase reaching overall lower cell densities and reduced maximum ANT titers of 1 ± 0.14 g/L (7.42 mM). Of all glucose/xylose ratios tested, the initially tested 1 % glucose / 3 % xylose mixture yielded the most ANT. Furthermore, *C. glutamicum* ANT2 harboring the isomerase pathway was also cultivated with 4 % glucose as the sole carbon source. Surprisingly, this strain produced up to 461.3 ± 22.7 mg/L (3.4 mM) ANT within 24 h, whereas the strain without expression of *xylA_Xc_* and *xylB_Cg_* accumulated only 370.4 ± 14.3 mg/L (2.7 mM) ANT. Since the plasmid pEKEx3-*xylA_Xc_*-*xylB_Cg_* had a positive effect on product formation, it was used in all subsequent experiments with glucose as the sole carbon source along with pMKEx2-*aroF*\**_EcCg_*-*tkt_Cg_*. Noteworthy, in addition to the formation of glycosyl- anthranilate in the presence of glucose, another peak with a shorter retention time and identical absorption properties as ANT emerged during HPLC analyses of all cultures with glucose/xylose mixtures. This peak could have been evoked by the presence of xylosyl-anthranilate, as it only occurred when xylose was used as a cosubstrate (Fig. S6).

Previously, a S38R-substitution was described in the ANS of *Brevibacterium lactofermentum* (*C. glutamicum* ssp. *lactofermentum*), which desensitizes this enzyme to intracellular TRP-concentrations of up to 10 mM (Matsui *et al*., 1987). With the aim to improve the flux towards ANT, the same point mutation was also introduced into *trpE* of *C. glutamicum* ANT2, yielding *C. glutamicum* ANT2 ANS-S38R (*C. glutamicum* ANT3). This variant was transformed with the respective plasmids enabling ANT synthesis and cultivated in shake flasks with 4 % glucose and 1 % glucose / 3 % xylose carbon source mixtures (Fig. 3). *C. glutamicum* ANT2 pMKEx2-*aroF*\**_EcCg_*-*tkt_Cg_* pEKEx3-*xylA_Xc_*_-_*xylB_Cg_* accumulated 355.3 ± 36.1 mg/L (2.59 mM) ANT from glucose within 24 h, while the ANT titer determined for the strain harboring ANS-S38R was almost doubled (624.2 ± 64.5 mg/L (4.55 mM) ANT). Using a mixture of glucose and xylose, *C. glutamicum* ANT3 pMKEx2-*aroF*\**_EcCg_*-*tkt_Cg_* pEKEx3-*xylA_Xc_*-*xylB_Cg_* reached a maximum product titer of 2.9 ± 0.39 g/L (21.04 mM) ANT, meaning that the product titer could be increased by almost 1 g/L compared to the variant with the wild-type ANS.

**Fig. 3:**
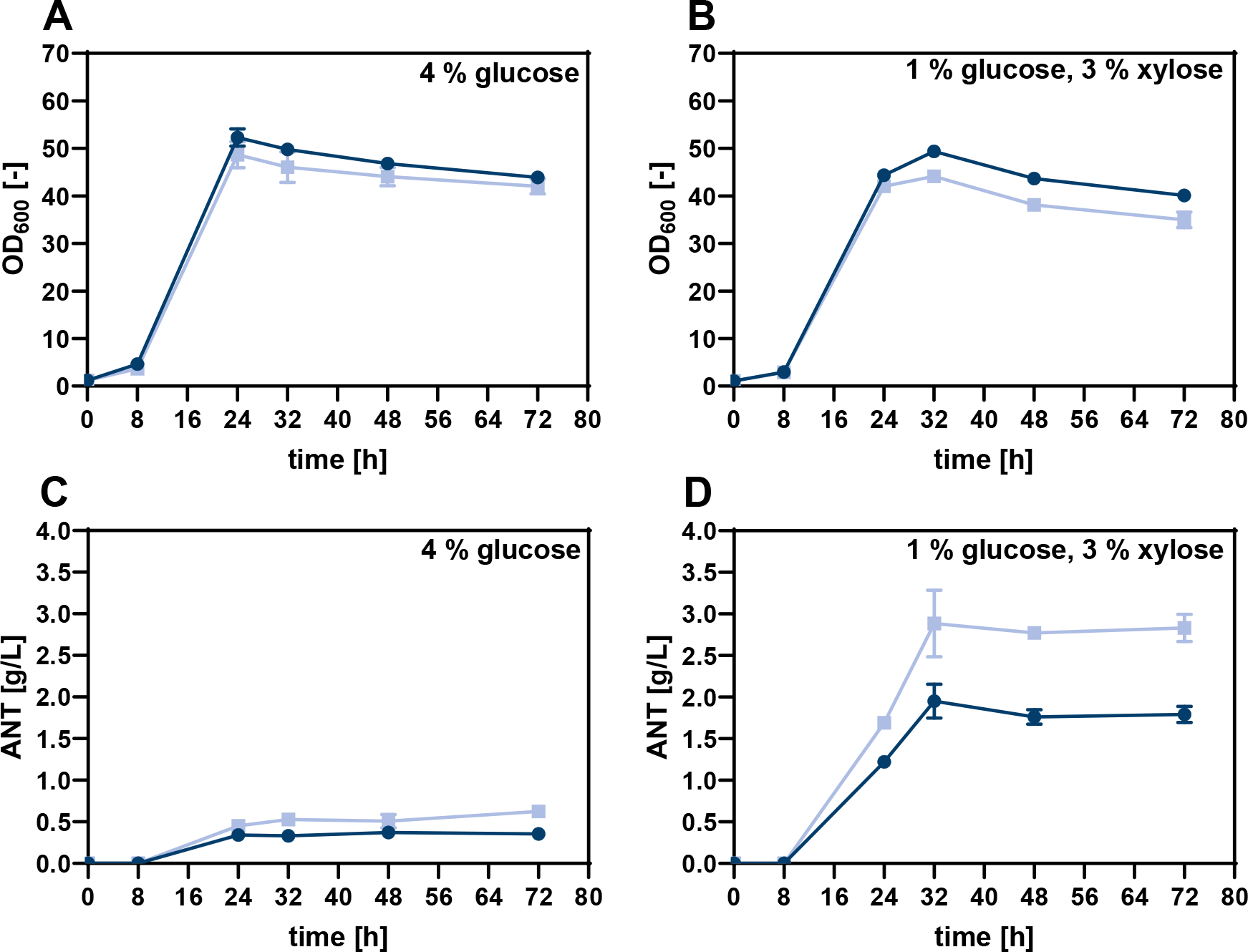
Growth and ANT production from glucose or glucose/xylose mixtures with *C. glutamicum* ANT3 bearing a feedback-resistant anthranilate synthase (ANS-S38R). (**A, B**). Growth (OD_600_) of the strains *C. glutamicum* ANT2 (control, circles) and *C. glutamicum* ANT3 (squares) with either 4 % glucose or a mixture of 1 % glucose / 3 % xylose as carbon source. (**C, D**) Determination of the ANT titer in the culture supernatants by HPLC. Both strains carry pMKEx2-*aroF*\**_EcCg_*-*tkt_Cg_* for ANT production and pEKEx3- *xylA_Xc_*-*xylB_Cg_* for xylose utilization via the isomerase pathway. The depicted data represent mean values and standard deviation of biological triplicates.

Noteworthy, a major by-product identified quantitatively by GC-TOF-MS analysis of the culture supernatants of the *C. glutamicum* ANT3 strains was SA, indicating that the ATP-dependent phosphorylation of SA to S3P might be a bottleneck within the SA pathway at this stage of metabolic engineering. In *C. glutamicum*, this step is catalyzed by SA kinase encoded by *aroK* (Syukur Purwanto *et al*., 2018). To reduce the accumulation of SA and thus increase the carbon flux through the SA pathway, the initial translation efficiency of *aroK* was increased by a start codon replacement in *aroK* (GTG→ATG). The resulting strain *C. glutamicum* ANT3 ATG-*aroK* (*C. glutamicum* ANT4) was transformed with the production plasmids and cultivated in shake flasks to investigate the effect of the start codon exchange on ANT and SA titer (Fig. S7). The cultivations with glucose as the sole carbon and energy source showed that the start codon replacement did not greatly affect the overall ANT titer, but could reduce the amount of SA accumulated within 24 h by more than 65 % (16.2 ± 2.6 mg/L (0.1 mM) SA vs. 46.5 ± 8.7 mg/L (0.3 mM) SA). However, towards the end of the cultivation, the same SA concentration of 10 mg/L (0.1 mM) could be determined in the supernatant of both strain variants. In contrast, the*C. glutamicum* ANT3 variant cultivated on a glucose/xylose mixture accumulated 374.7 ± 6.6 mg/L (2.2 mM) SA within 24 h. Here the *aroK* start codon replacement could reduce the SA concentration only by 14.6 % to 320.6 ± 20.4 mg/L (1.8 mM) SA.

### 3.4 Biosensor-guided engineering of the anthranilate synthase of *C. glutamicum*

The introduction of the feedback-relieved ANS-S38R–variant proved to be a key modification for high ANT production with *C. glutamicum* in this study. However, with the aim to find out whether the feedback inhibition could be further reduced, we set out to identify additional amino acid residues in the ANS, which are essential for TRP-mediated feedback inhibition. The ANS from *Salmonella typhimurium* shows 47.9 % homology to the enzyme of *C. glutamicum* and a previous study indicated that amino acid residues E39, S40, M293, F294, and C465 of this enzyme are associated with feedback-inhibition by TRP (Morollo & Eck, 2001). Since four residues are also conserved in the ANS from *C. glutamicum* (E37, S38, M285, and C461), improved ANS variants with an increased TRP-feedback resistance might be obtainable.

In order to access such ANS variants, the codons for the residues E37, S38, M285, and C461 of an episomally encoded *trpE*-gene of *C. glutamicum* were targeted by SSM, and the resulting libraries were subjected to a biosensor-based *in vivo*-screening in MTPs. Prior to these experiments, the genome-encoded *trpE* was deleted for screening purposes to avoid cross-talk with the native ANS. This *trpE*-deficient strain required TRP-supplementation in the defined CGXII medium but episomal *trpE* expression could restore growth (Fig. S8). Subsequently, four individual SSM libraries of *trpE* encoded on a pJC1-plasmid were generated using NNK- degenerated oligonucleotides. Noteworthy, before transformation of the pJC1-plasmids carrying the mutated ANS-variants, *aroF*\**_EcCg_* was integrated into the intergenic region (IGR09) between cg0432 and cg0435 in the genome of *C. glutamicum*, since the pJC1- and the pMKEx2-plasmids are incompatible. The resulting *C. glutamicum* ANT4 IGR9::P*dapA*(A16)-*aroF*\**_EcCg_* (*C. glutamicum* ANT5) variant allowed for the formation of 380 ± 17.1 mg/L (2.8 mM) ANT from glucose within 24 h. Hence this strain is inferior to the *C. glutamicum* ANT4 strain enabling plasmid-based expression of *aroF*\**_EcCg_* and *tkt*, but suitable for the envisioned ANS-screening (for details of the conducted strain characterization, the reader is referred to the supplementary material, Fig. S9). After transformation, 150 variants for each library, corresponding to a theoretical library completeness of 95.5 % (GLUE-IT algorithm (Firth & Patrick, 2008)), were screened.

The pSenCA biosensor for the intracellular detection of cinnamic acid and phenylpropionic acid in *E. coli* was previously engineered to also detect other small aromatic molecules of biotechnological interest (Flachbart *et al*., 2021, 2019). One of these customized biosensor variants is pSen6MSA, engineered for the detection of 6-methylsalicylic acid (6MSA). Due to the structural similarity of ANT and 6MSA (Fig. 1, 4A), we assumed that this biosensor might also recognize ANT, which would make an *E. coli* strain bearing this sensor a screening tool for the detection of ANT-accumulating *C. glutamicum* variants. To test the applicability of pSen6MSA for this purpose, *E. coli* DH10B Δ*hcaREFCBD* pSen6MSA was cultivated in the presence of 0-20 mM ANT or *C. glutamicum* culture supernatants containing 0-2 mM microbially produced ANT (Fig. S10, S11). ANT could indeed induce the biosensor and the optimum inducer concentration was determined to be 0.6 mM ANT. Higher ANT concentrations resulted in impaired growth of the *E. coli* strain restricting the overall fluorescence. Since up to 1.8 mM SA typically accumulate in supernatants of ANT-producing *C. glutamicum* cultures, it was further examined if SA can also act as an inducer of the pSen6MSA. However, no specific fluorescence was observed upon the addition of 0-20 mM SA, indicating that no cross-talk was to be expected (Fig. S12).

For screening, all 600 clones of the four site-saturated *C. glutamicum* ANT5 Δ*trpE* libraries were individually cultivated in MTPs for ANT production. Subsequently, culture supernatants were transferred to a second MTP with cultures of the pSen6MSA bearing *E. coli* sensor strain for the semi-quantitative detection of ANT. For a detailed analysis, ANT concentrations in the supernatant of the controls and of three clones with lower, equal, or higher specific fluorescence of each library screened were determined by HPLC (Tab. S3-6). In the case of the ANS-E37x, only a few variants showed an increased fluorescence response, but the DNA-sequencing of these clones only returned wild-type *trpE* sequences (Fig. S13). Screening of the ANS-M285x library yielded a few variants with an increased biosensor response (Fig. S15). However, a detailed HPLC analysis of the supernatants indicated an equal or lower ANT titer so DNA sequencing was omitted (Tab. S5). Most clones reached the same fluorescence as the control, which suggested that ANS-M285 might not be important for feedback inhibition. The biosensor-based screening of the ANS-S38x library identified three clones with an increased fluorescence signal (Fig. S14). Interestingly, DNA sequencing revealed S38A and S38G substitutions, and the already known S38R substitution could not be retrieved. Finally, screening of the ANS-C461x library yielded 15 variants with an increased biosensor response (Fig. 4B). DNA-sequencing revealed that all *trpE*-variants were characterized by mutations leading to a C461G substitution, suggesting that this amino acid residue is indeed important for feedback inhibition.

**Fig. 4:**
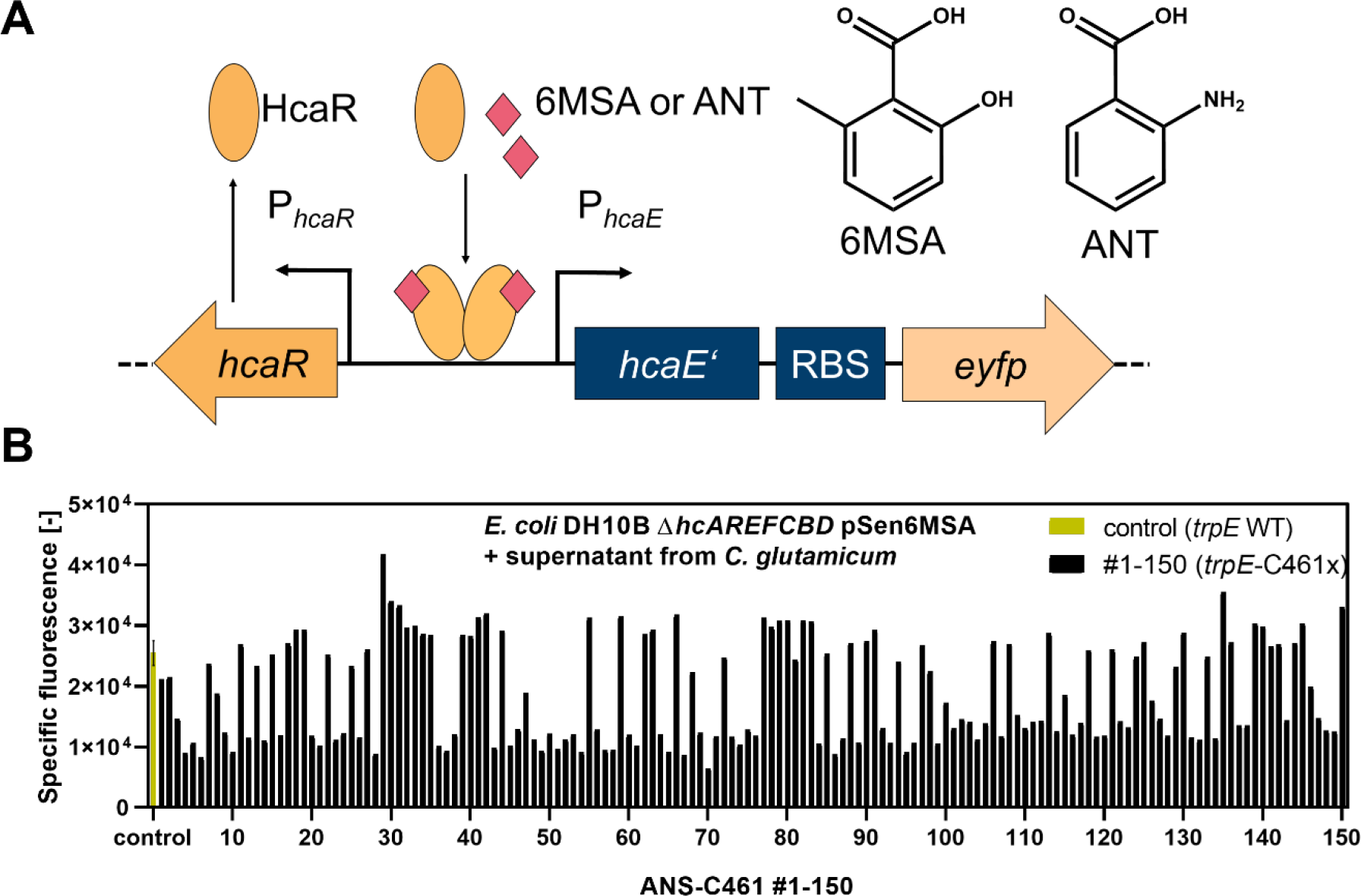
Biosensor-based screening of anthranilate synthase libraries. (**A**) Schematic of the pSen6MSA biosensor, 6MSA, and ANT structure to demonstrate the chemical similarity between both molecules. (**B**) Characterization of a *C. glutamicum* strain library with randomly mutagenized nucleotide triplet encoding ANS-C461.

Subsequently, the two mutations leading to the two amino acid substitutions ANS-S38A and ANS-S38G were individually introduced into the genome of *C. glutamicum* ANT5 to confirm the positive effects on ANT production in shake flask cultivations in comparison to the previously introduced ANS-S38R substitution (Fig. S16). Growth of all strain variants was indistinguishable and all variants reached a similar ANT titer of 2.7-2.9 g/L (19.7-20.8 mM) after 32 h. Subsequently, the mutation ANS-C461G was combined with the previously introduced ANS- S38R/A/G mutation to identify any synergistic effects of pairwise *trpE*-mutation on product formation (Fig. 5). All strains reached a final OD_600_ of 45-50 after 24 h (Fig 5A). For all ANS double mutants, a slightly increased ANT titer could be determined in comparison to the respective S38X variant after 48 h. Although this effect was neglectable at the end of each cultivation, the observed faster product formation hints indeed towards a combinatorial effect of both mutations in all mutated ANSs (Fig. 5 B-D).

**Fig. 5:**
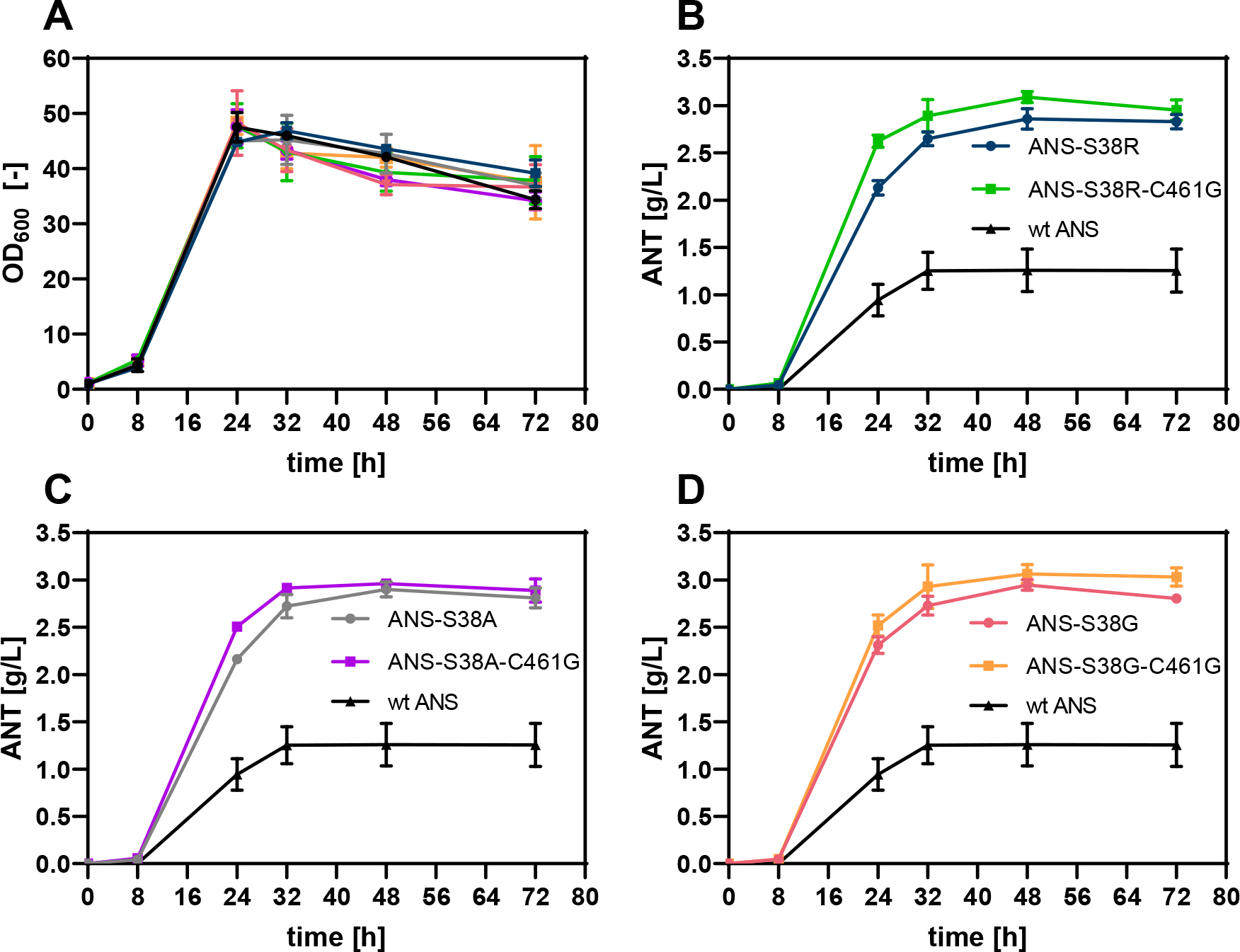
Growth and ANT production of different *C. glutamicum* strains with mutated anthranilate synthases. (**A**) Growth (OD_600_) and ANT titer of *C. glutamicum* ANT5 pMKEx2-*aroF*\**_EcCg_*-*tkt_Cg_* pEKEx3- *xylA_Xc_*-*xylB_Cg_* harboring (**B**) ANS-S38R or ANS-S38R-C461G, (**C**) ANS-S38A or ANS-S38A-C461G or (**D**) ANS-S38G or ANS-S38G-C461G. The depicted data represent mean values and standard deviation of biological triplicates.

### 3.5 Laboratory-scale production of anthranilate

For laboratory-scale production of ANT, *C. glutamicum* ANT5 ANS-C461G (*C. glutamicum* ANT6) was cultivated in pH-controlled bioreactors using defined CGXII medium (without urea and MOPS buffer) in batch and fed-batch mode (Fig. 6). In the batch mode, *C. glutamicum* ANT6 reached a maximum OD_600_ of 53.9 ± 0.9 after 23.3 h, before the optical density of the culture dropped slightly over the course of the cultivation. ANT production set in after 12 h of cultivation time and was finished after 27.9 h with a maximum product titer of 2.6 ± 0.02 g/L (19.1 mM). Modelling of the resulting data indicated a diauxic behavior of *C. glutamicum* ANT6, first consuming glucose exclusively for biomass formation, followed by xylose utilization solely for ANT production (Tab. S7). The low product yield of only 0.06 g_ANT_ g ^-1^ can be explained by the accumulation of side products, predominantly SA.

**Fig. 6:**
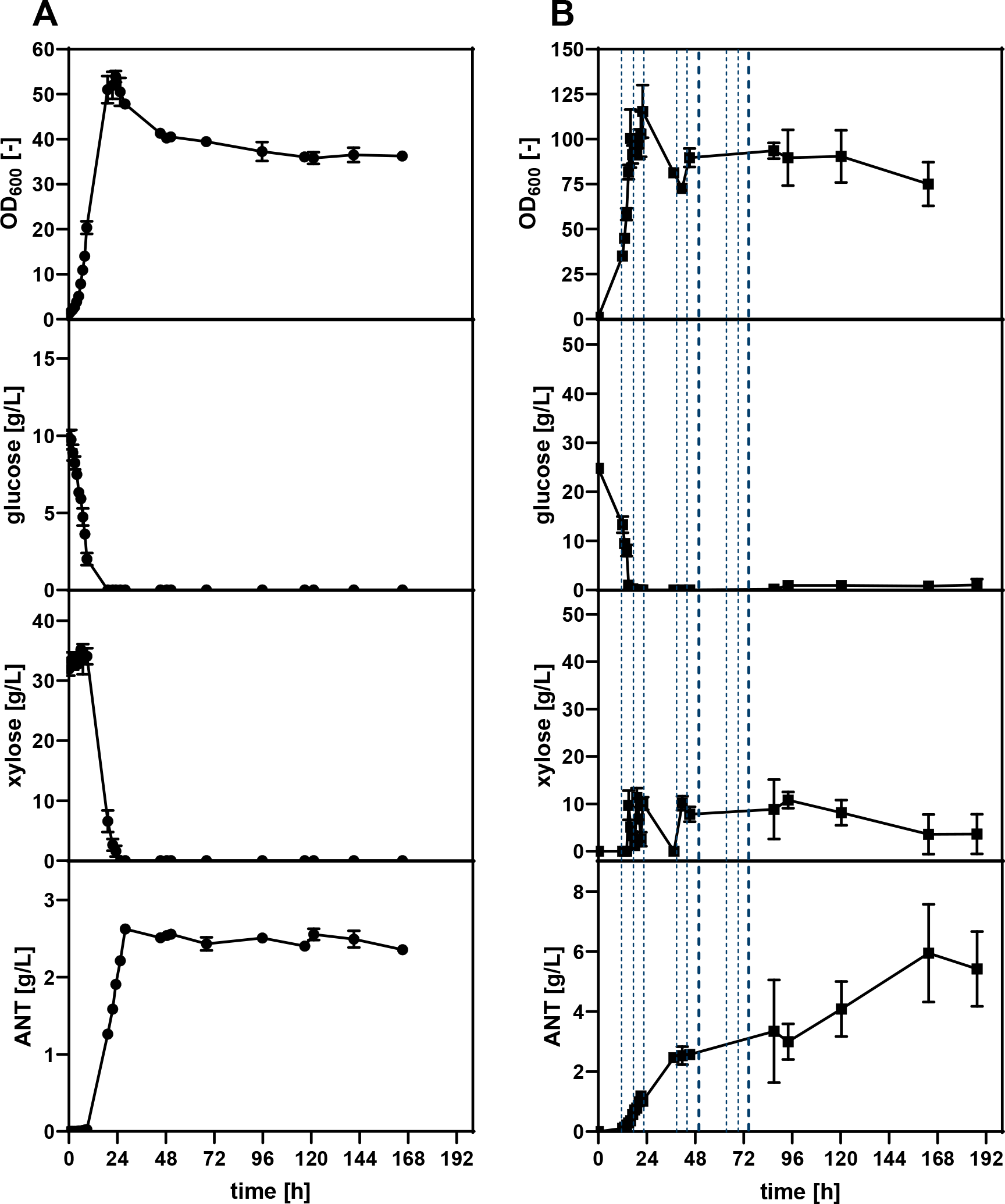
**Batch and pulse-fed**-**batch bioreactor cultivation of *C. glutamicum* ANT6 for ANT production.** Comparison of growth (OD_600_), glucose/xylose consumption, and ANT production in bioreactor cultivations of the strain *C. glutamicum* ANT6 harboring the expression plasmids pMKEx2-*aroF*\**_EcCg_*-*tkt_Cg_* and pEKEx3- *xylA_Xc_*-*xylB_Cg_* in (**A**) batch (circles) and (**B**) fed-batch mode (squares). In fed-batch cultivations, 10 g xylose (67 mM, dashed lines) or 20 g xylose (133 mM, thick dashed lines) were pulsed at defined time points. The depicted data represent mean values and standard deviation of biological duplicates.

During fed-batch mode, *C. glutamicum* ANT6 was first cultivated with glucose in batch cultivation for initial biomass formation, followed by a constant glucose feed and xylose pulses at defined time points. Again, the strain showed the diauxic behavior already observed during batch mode cultivation, but product formation of *C. glutamicum* ANT6 continued after the last xylose pulse. The strain reached an OD_600_ of 115.4 ±10.4 after 22 h and a maximum ANT titer of 5.9 ± 1.4 g/L (43.3 mM) after 168 h.

*C. glutamicum* ANT6 requires additional engineering to further reduce by-product accumulation, but the results obtained highlight the potential of the constructed *C. glutamicum* ANT variants for the production of ANT and other aromatic compounds from xylose containing feedstocks.

## 4 Discussion

In this study, the microbial production of ANT with *C. glutamicum* was established, employing rational and semi-rational engineering strategies. Initial experiments on product toxicity were performed due to known antimicrobial properties of ANT (Csonka, 1989; Li *et al*., 2017). However, only a minor effect of ANT on the growth of *C. glutamicum* was observed, making this organism a more suitable host for the production of ANT than *P. putida*, which cannot grow at ANT concentrations exceeding 10 g/L (72.92 mM) (Kuepper *et al*., 2015).

In order to avoid the formation of possible by-products, the genes *nagD* and *qsuD* were deleted, whose gene products are involved in the synthesis of glycerol and QA (Jojima *et al*., 2012; Teramoto *et al*., 2009). Single deletion of *nagD* or *qsuD* had no impact on ANT production, only the double deletion increased the product titer significantly. Deletion of *nagD* might lead to increased availability of NADPH, which in turn could be used by *qsuD*-encoded NADPH-dependent QA/SA dehydrogenase to form QA, resulting in no increased ANT titer (Jojima *et al*., 2012). Glycerol and QA both could not be detected in the culture supernatant by HPLC although the formation of glycerol has been already described during SA-synthesis with *C. glutamicum* (Kogure *et al*., 2016). However, in the context of this particular study, glycerol accumulation was only detectable during cultivations in which SA titers exceeding 100 g/L were obtained. Hence, glycerol and QA concentrations present in the culture supernatants of the *C. glutamicum* strains constructed in this study might have been below the detection limit.

Initially, low ANT titers obtained in cultivations with glucose as sole carbon and energy source, could be markedly increased when using glucose/xylose mixtures since xylose is rapidly converted to E4P via the isomerase pathway and reactions of the PPP (Radek *et al*., 2014). This in turn increased the carbon flux through the SA pathway, which was also evident from the observed accumulation of the central SA pathway intermediate SA. Utilization of this mixed carbon substrate allowed for the production of ANT at g/L scale, outperforming ANT synthesis with *P. putida* (1.5 g/L (10 mM) ANT) (Kuepper *et al*., 2015). The maximum ANT titer reached in pulsed-fed-batch bioreactors with *C. glutamicum* was 5.9 ± 1.4 g/L (43.3 mM), which exceeded the ANT titer determined for *B. subtilis* (3.5 g/L (26 mM)), but was lower compared to previous experiments with *E. coli* (14 g/L (102 mM)) (Balderas-Hernández *et al*., 2009; Cooper *et al*., 1995). However, these high product titers were achieved in complex media limiting the comparability of the obtained results. Noteworthy, we observed an increased product formation in cultivations with glucose as the sole carbon source when genes for the heterologous isomerase pathway for xylose utilization were episomally expressed. In general, xylose isomerases are known to have a rather broad substrate spectrum also accepting glucose as a substrate, which is isomerized to fructose (Blow *et al*., 1992; Tsumura & Sato, 1965). Subsequently, fructose is exported and re-imported by the fructose-specific phosphotransferase system yielding fructose- 1-phosphate, which then enters glycolysis after phosphorylation to fructose-1,6-bisphosphate (Dominguez & Lindley, 1996; Ikeda, 2012). Thus, the xylose isomerase side activity might increase glucose utilization via glycolysis in the performed cultivations.

During previous engineering of *E. coli*, *P. putida* and *C. glutamicum* for ANT overproduction, *trpD* was deleted to increase ANT accumulation (Balderas-Hernández *et al*., 2009; Kuepper *et al*., 2015; Luo *et al*., 2019). However, we favored the replacement of the ATG start codon of *trpD* by GTG to reduce the translation efficiency to avoid any requirement for TRP supplementation, which is inevitable when deleting this essential gene.

As was the case with the DAHP synthase, removing the feedback inhibition of the key enzyme ANS was one of the decisive factors to improve product formation. A similar strategy was pursued regarding the microbial synthesis of ANT with *P. putida*, expressing *trpE^S40^G* from *E. coli*, which was achieved via plasmid-based expression (Kuepper *et al*., 2015). The performed biosensor-based screening of focused libraries for ANS variants with further reduced feedback-inhibition showed that the obtained S38A/G substitutions had the same effect as the previously described S38R substitution, which confirms the assumption that the hydroxy group of the native SER residue is essential for TRP binding (Matsui *et al*., 1987). The other amino acid substitution leading to relieved TRP-mediated feedback inhibition was C461G. In combination with S38A or S38G, this novel amino acid substitution had a positive additive effect on ANT accumulation, indicating that the degree of feedback-resistance can be gradually adjusted by substitutions of more than one key residue.

Batch and pulsed-fed-batch cultivations of *C. glutamicum* ANT6 showed an interesting diauxic growth/production behavior during which glucose was first metabolized and used for biomass formation. Subsequently, xylose was consumed and channeled towards ANT production and by-product formation. Noteworthy, *C. glutamicum* is known for its ability to cometabolize glucose with acetate, lactate or fructose among others (Cocaign *et al*., 1993; Dominguez *et al*., 1998, 1997). The only known examples of diauxic substrate consumption of *C. glutamicum* are the sequential metabolism of glucose before glutamate and glucose before ethanol (Arndt *et al*., 2008; Krämer *et al*., 1990). However, in case of the engineered catabolization of the xylose via the isomerase pathway, a preferential utilization of glucose over xylose has been already observed several times in different *C. glutamicum* species (Kawaguchi *et al*., 2006; Radek *et al*., 2014). A possible reason for this could be a side activity of the xylose isomerases already discussed in the context of the improved ANT production in cultivations with glucose when the genes for the isomerase pathway were episomally expressed. In the case of the preferential glucose utilization observed in the bioreactors, a competitive competition of glucose and xylose for the binding sites of the xylose isomerases might delay an efficient xylose utilization.

Taken together, by combining rational strain engineering and biosensor-guided semi-rational enzyme engineering approaches, an attractive *C. glutamicum* strain for ANT production from glucose/xylose mixtures could be constructed. This variant represents a promising starting point for the construction of other production strains for a broader spectrum of aromatic compounds of commercial interest.

## Associated content

### Supplementary information

Supplementary information accompanies this paper. Additional information contains a list of constructed strains and plasmids, oligonucleotides used, HPLC chromatograms displaying peaks of anthranilate, *N*- glycosyl-anthranilate and *N*-xylosyl-anthranilate and additional cultivation results.

### Author information Corresponding author

Jan Marienhagen - Institute of Bio- and Geosciences, IBG-1: Biotechnology, Forschungszentrum Jülich, D- 52062 Jülich, Germany; https://orcid.org/0000-0001-5513-3730; Phone: +49 2461 61 2843; Fax: +49 2461 61 2710

### Authors

Mario Mutz - Institute of Bio- and Geosciences, IBG-1: Biotechnology, Forschungszentrum Jülich, D-52425 Jülich, Germany; Institute of Biotechnology, RWTH Aachen University, Worringer Weg 3, D-52074 Aachen, Germany; https://orcid.org/0000-0003-1716-6931

Vincent Brüning **-** Institute of Bio- and Geosciences, IBG-1: Biotechnology, Forschungszentrum Jülich, D- 52425 Jülich, Germany; https://orcid.org/0000-0002-8449-8904

Christian Brüsseler - Institute of Bio- and Geosciences, IBG-1: Biotechnology, Forschungszentrum Jülich, D-52425 Jülich, Germany; https://orcid.org/0000-0003-028-7827

Moritz-Fabian Müller - Institute of Bio- and Geosciences, IBG-1: Biotechnology, Forschungszentrum Jülich, D-52425 Jülich, Germany; https://orcid.org/0000-0002-2503-2768

Stephan Noack - Institute of Bio- and Geosciences, IBG-1: Biotechnology, Forschungszentrum Jülich, D- 52425 Jülich, Germany; https://orcid.org/0000-0001-9784-3626

### Declaration of competing interests

The authors declare that they have no conflict of interest.

### Author contributions

CB and JM conceived the study. MM and VB performed the experiments. MFM and SN performed the bioreactor cultivations for laboratory-scale production of anthranilate. JM supervised the study. MM prepared the figures and wrote the manuscript with input from all authors.

## Acknowledgements

Not applicable.

## Funding

JM acknowledges funding from the European Research Council (ERC) under the European Union’s Horizon 2020 research and innovation program under grant agreement no 638718. JM and MM acknowledge funding from the European Union’s Horizon 2020 research and innovation program under grant agreement no. 953073 (UPLIFT).

## Data availability

The data used in this study can be made available upon request to the corresponding author.

## Notes

### Competing Interest Statement

The authors have declared no competing interest.

