## Supplementary Material for "Metabolic engineering of *Corynebacterium glutamicum* for the production of anthranilate from glucose and xylose"

| Strain or plasmid | Characteristics | Source/reference |
|---|---|---|
| ***C. glutamicum*** **strains** | | |
| DelAro[5] $P_{O6}$-*iolT1* | Prophage-free derivative of ATCC 13032 with an in-frame deletion of cg0344-cg0347, cg2625-cg2640, cg1226, cg502, and cg3349-cg3354 (DelAro[5]). Two point mutations in the promotor of the inositol transporter gene *iolT1* abolish repression of *iolT1* by IolR | (Kallscheuer & Marienhagen, 2018) |
| DelAro[5] $P_{O6}$-*iolT1* GTG-*trpD* (ANT1) | DelAro[5] $P_{O6}$-*iolT1* derivative with the start-codon exchange of *trpD* (ATG→GTG) | This work |
| ANT1 Δ*nagD* | ANT1 derivative with an in-frame deletion of cg2474 | This work |
| ANT1 Δ*qsuD* | ANT1 derivative with an in-frame deletion of cg0504 | This work |
| ANT1 Δ*nagD* Δ*qsuD* (ANT2) | ANT1 derivative with an in-frame deletion of cg2474 and cg0504 | This work |
| ANT2 ANS-S38R (ANT3) | ANT2 derivative harboring a mutated anthranilate synthase carrying the amino acid substitution S38R | This work |
| ANT3 ATG-*aroK* (ANT4) | ANT3 derivative with a start-codon exchange of *aroK* (GTG → ATG) | This work |
| ANT4 IGR9::*aroF*$^*_{EcCg}$ (ANT5) | ANT4 derivative with genomic integration of *aroF*$^*_{EcCg}$ encoding AroF* from *E. coli* (codon-optimized gene) under control of the *dapA* promotor variant A16. | This work |
| ANT5 Δ*trpE* | ANT5 derivative with an in-frame deletion of cg3359 | This work |
| ANT5 ANS-C461G | ANT5 derivative harboring a mutated anthranilate synthase carrying the amino acid substitution S38R and C461G | This work |
| ANT5 ANS-S38G-C461G | ANT5 derivative harboring a mutated anthranilate synthase carrying the amino acid substitution S38G and C461G | This work |

| | | |
|---|---|---|
| ANT5 ANS-S38A-C461G | ANT5 derivative harboring a mutated anthranilate synthase carrying the amino acid substitution S38A and C461G | This work |
| ANT5 ANS-C461G Δ*pyk* | ANT5 ANS-C461G derivative with an in-frame deletion of cg2291 | This work |
| ***E. coli*** strains | | |
| *E. coli* DH5α | $F^-$Φ80*lacZ*ΔM15 Δ(*lacZYA- argF*)U169 *recA1 endA1 hsdR17* (rK-, mK+) *phoA supE44λ- thi-1 gyrA96 relA1* | Invitrogen (Karlsruhe, Deutschland) |
| *E. coli* TOP10 | $F^-$*mcr*A Δ(*mrr-hsd*RMS-*mcr*BC) φ80*lac*ZΔM15 Δ*lac*X74 *rec*A1 *ara*D139 Δ(*ara-leu*) 7697 *gal*U *gal*K *rps*L ($Str^R$) *end*A1 *nup*G $λ^-$ | Thermo Fisher Scientific (Waltham, MA, USA) |
| *E. coli* DH10B Δ*hcaREFCBD* | $F^-$*mcr*A Δ(*mrr-hsd*RMS-*mcr*BC) φ80*lac*ZΔM15 Δ*lac*X74 *rec*A1 endA1 *ara*D139 Δ(*ara*, *leu*)7697 *gal*U *gal*K $λ^-$*rps*L *nup*G Δ*hcaREFCBD* | (Flachbart *et al.*, 2021) |
| **Plasmids** | | |
| pMKEx2 | *kan*$^r$; *E. coli*-*C. glutamicum* shuttle vector (*lacI*, $P_{T7}$, lacO1, pHM1519 $ori_{Cg}$; ACYC177 $ori_{Ec}$) | (Kortmann *et al.*, 2015) |
| pMKEx2-$aroF^*_{EcCg}$-$tkt_{Cg}$ | pMKEx2 derivative for expression of $aroF^*_{EcCg}$ (codon-optimized gene) encoding feedback-resistant DAHP synthase from *E. coli* and $tkt_{Cg}$ encoding transketolase from *C. glutamicum*, whose native TTG start codon was exchanged by ATG. | This work |
| pEKEx3 | *spec*$^r$; *E. coli*-*C. glutamicum* shuttle vector (*lacI*, $P_{tac}$, lacO1, pBL1$ori_{Cg}$; pUC$ori_{Ec}$) | (Gande *et al.*, 2007) |
| pEKEx3-$xylA_{Xc}$-$xylB_{Cg}$ | pEKEx3 derivative for expression of $xylA_{Xc}$ (XCC1758) encoding xylose isomerase from *Xanthomonas campestris*) and $xylB_{Cg}$ encoding xylulokinase from *C. glutamicum* | (Meiswinkel *et al.*, 2013) |
| pK19*mobsacB* | *kan*$^r$; vector for allelic exchange in *C. glutamicum* (pK18 oriVEc *sacB* lacZα) | (Schäfer *et al.*, 1994) |
| pK19mobsacB-GTG-*trpD* | pK19*mobsacB* derivative for the start-codon exchange of *trpD* (GTG→ATG) | This work |

| | | |
|---|---|---|
| pK19*mobsacB*-Δ*nagD* | pK19*mobsacB* derivative for the in-frame deletion of *nagD* | This work |
| pK19*mobsacB*-Δ*qsuD* | pK19*mobsacB* derivative for the in-frame deletion of *qsuD* | This work |
| pK19*mobsacB*-ANS-S38R | pK19*mobsacB* derivative for site-directed mutagenesis of *trpE* encoding anthranilate synthase component I resulting in the amino acid substitution S38R. | This work |
| pK19*mobsacB*-ATG-*aroK* | pK19*mobsacB* derivative for the start-codon exchange of *aroK* (GTG→ATG) | This work |
| pK19*mobsacB*-IGR9 | pK19*mobsacB* derivative for genomic integration between cg0432 and cg0435 | This work |
| pK19*mobsacB*-IGR9::P*dapA*-A16-$aroF^{*}_{EcCg}$ | pK19*mobsacB*-IGR9 derivative for the genomic integration of $aroF^{*}_{EcCg}$, whose gene expression is under control of the *dapA* promotor variant A16 | This work |
| pK19*mobsacB*-Δ*pyk* | pK19mobsacB derivative for in-frame deletion of *pyk* | (Labib *et al.*, 2021) |
| pK19*mobsacB*-ANS-S38A | pK19*mobsacB* derivative for site-directed mutagenesis of *trpE* encoding anthranilate synthase component I resulting in the amino acid substitution S38A. | This work |
| pK19*mobsacB*-ANS-S38G | pK19*mobsacB* derivative for site-directed mutagenesis of *trpE* encoding anthranilate synthase component I resulting in the amino acid substitution S38G | This work |
| pK19*mobsacB*-ANS-C461G | pK19*mobsacB* derivative for site-directed mutagenesis of *trpE* encoding anthranilate synthase component I resulting in the amino acid substitution C461G | This work |
| pSen6MSA | Episomal transcriptional biosensor inducing *eyfp* expression in response to the presence of 6-methylsalicylic acid or ANT, $Kan^{R}$ | (Flachbart *et al.*, 2021) |
| pJC1 | $Kan^{r}$; *E. coli*/*C. glutamicum* shuttle ($oriV_{E. coli}$, $oriV_{C. glutamicum}$) | (Cremer *et al.*, 1990) |
| pJC1-*trpE* | pJC1 derivative for expression of $trpE_{Cg}$. Gene expression is under the control of the *dapA* promotor variant A16 | This work |

**Tab. S2: Oligonucleotides used in this study**

| Oligonucleotide | Sequence [5`→3`] |
|---|---|
| *aroF**$_{EcCg}$ fwd | CCAAGCGTGTTGAAGATGGTGGGGAAT |
| *aroF**$_{EcCg}$ rev | CCAAGCGTGTTGAAGATGGTGGGGAAT |
| *tkt*$_{Cg}$ fwd | CCAAGCGTGTTGAAGATGGTGGGGAAT |
| *tkt*$_{Cg}$ rev | CCAAGCGTGTTGAAGATGGTGGGGAAT |
| GTG-*trpD* Up fwd | ATCCCCGGGTACCGAGCTCGCCTTTCACCTGGACCTGG |
| GTG-*trpD* Up rev | GAATCAAATCCTTTTTTTATTAGTTCGC |
| GTG-*trpD* Down fwd | ATAAAAAAAGGATTTGATTCGTGACTTCTCCAGCAACACTG |
| GTG-*trpD* Down rev | TTGTAAAACGACGGCCAGTGCTGCACATGCGCAATCGC |
| GTG-*trpD* Check fwd | GCCAGTGGAAACCATTTTGGCAGCC |
| GTG-*trpD* Check rev | CCAAGCGTGTTGAAGATGGTGGGGAAT |
| Δ*nagD* Up fwd | TGCATGCCTGCAGGTCGACTTTTAGACCCGGGGTACGG |
| Δ*nagD* Up rev | GTACGTGAAATGAAATGTTCACTGTCATAACACC |
| Δ*nagD* Down fwd | GAACATTTCATTTCACGTACCAGATGAGCAGC |
| Δ*nagD* Down rev | TTGTAAAACGACGGCCAGTGCTCGGGGAAGCCAGGTGA |
| Δ*nagD* Check fwd | CGAGGGTGAAGCCGTGGATGAACAC |
| Δ*nagD* Check rev | GTGTTCTTCGACTGCTTCGCCGTAG |
| Δ*qsuD* Up fwd | TGCATGCCTGCAGGTCGACTCTAGCATCCCGAACTAGC |
| Δ*qsuD* Up rev | TGCGGGAGACGAGAATACTGTCGTTCATATTTTTG |
| Δ*qsuD* Down fwd | CAGTATTCTCGTCTCCCGCATGCGGGAA |
| Δ*qsuD* Down rev | TTGTAAAACGACGGCCAGTGGACCAAACTAGGTGACCGAAAACT TTTTC |
| Δ*qsuD* Check fwd | GTTTTGCAGGTTGGGGCGAAACAAACG |
| Δ*qsuD* Check rev | GCGTTTTAGGGGCTTAGACGCGATTCT |
| ATG-*aroK* Up fwd | TGCATGCCTGCAGGTCGACTTTCTGGCGAAGATCGCCTC |
| ATG-*aroK* Up rev | GATCATTCATTTCATTACGCTCCATCCTCTAAAC |
| ATG-*aroK* Down fwd | GCGTAATGAAATGAATGATCAAATTCACTTAGATCATC |
| ATG-*aroK* Down rev | GCTCGGTACCCGGGGATCCTTTGGTGCGAAGTCGGTAG |
| ATG-*aroK* Check fwd | TGAGGCCGGAACCAATGTGGACATC |
| ATG-*aroK* Check rev | CGTGAAACCTTGGGCGTGGAAGTGC |
| Δ*trpD* Up fwd | ATCCCCGGGTACCGAGCTCGAATTCTTACCCTGCCGATGCGGG |
| Δ*trpD* Up rev | AGTAATCGATTGTTGCTGGAGAAGTCATGAATCAAATC |
| Δ*trpD* Down fwd | TCCAGCAACAATCGATTACTCAGAAAAGGAGTCTTCCAATGACTAGTAATAA TC |
| Δ*trpD* Down rev | TTGTAAAACGACGGCCAGTGAATTCCGGCAGCGAGTGCTGCGT |
| Δ*trpD* Check fwd | CAAGGCGTATGCCGTGTTGAATGCCATTGC |
| Δ*trpD* Check rev | AAAGACTTTGCCTTCACAGCGGCCGGCACCGTAGC |
| ANS-S38R Up fwd | ATCCCCGGGTACCGAGCTCGCGATTATCAACAAGACTCCGC |
| ANS-S38R Up rev | TATCAGCGCGTTCCAACAGGGCTGCATC |
| ANS-S38R Down fwd | CCTGTTGGAACGCGCTGATATCACCACC |
| ANS-S38R Down rev | TTGTAAAACGACGGCCAGTGAGCAATGAAAGCTTGTTGAGC |
| IGR9-P*dapA*-A16 fwd | TGCCGTCGGGAGGTATTTGCTTGGGCGATTGTTATGCAAAAG |
| IGR9-P*dapA*-A16 rev | GTTCAATTATACCTTCTTCCTTCATTTG |
| *aroF**$_{EcCg}$ Int. fwd | GGAAGAAGGTATAATTGAACCATATGACCGATGAACAGGTGCTG |
| *aroF**$_{EcCg}$ Int. rev | TCCCGGGATCTTAAGTGAGCTCTAGATTATGCCACGCGTGCGGT |
| IGR9-P*dapA*-A16-*aroF**$_{EcCg}$ Check fwd | TGCTGGCAGAATTCTCCTAATCCGGC |

| | |
|---|---|
| IGR9-P*dapA*-A16-$aroF^{*}_{EcCg}$ Check rev | AGTGGACAGATATTCTTCGAGATCG |
| PJC1-P*dapA*(A16)-fwd | CGACGCCGCAGGGGGATCCTTGCTTGGGCGATTGTTATG |
| PJC1-P*dapA*(A16)-rev | TCGTGCTCATATGGTTCAATTATACCTTCTTCC |
| PJC1-*trpE*-fwd | ATTGAACCATATGAGCACGAATCCCCATG |
| PJC1-*trpE*-rev | GCCATTGCTGCAGGTCGACTTCATCGGATGACCTCCAAAG |
| SSM-ANS-Glu-37 fwd | AGCCCTGTTGNNKAGCGCTGATATCAC |
| SSM-ANS-Glu-37 rev | GCATCATCTGCGGTTGTGCCAC |
| SSM-ANS-Ser-38 fwd | CTGTTGGAANNKGCTGATATCACCAC |
| SSM-ANS-Ser-38 rev | GGCTGCATCATCTGCGGTTGTGC |
| SSM-ANS-Met-285 fwd | CGTCGCCGTACNNKTTCTATATCCG |
| SSM-ANS-Met-285 rev | GGTTGGTGGCACGCAGCTGCAGATAAGC |
| SSM-ANS-Cys-461 fwd | GGCGATATGGATAATNNKATTGTTATTCGTTCGGCG |
| SSM-ANS-Cys-461 rev | ATTGCCGCGCAGGTACCCCACTG |
| ANS-S38x Up fwd | ATCCCCGGGTACCGAGCTCGCGATTATCAACAAGACTCCGC |
| ANS-S38x Up rev | TTCCAACAGGGCTGCATC |
| ANS-S38A Down fwd | ATGATGCAGCCCTGTTGGAAGCGGCTGATATCACCACCAAG |
| ANS-S38G Down fwd | ATGATGCAGCCCTGTTGGAAGGGGCTGATATCACCACCAAG |
| ANS-S38x Down rev | TTGTAAAACGACGGCCAGTGATAAGCAATGAAAGCTTGTTG |
| *trp* locus check primer II | GGTTTGGTAGGCGTGTTCGGTTGCG |
| ANS-C461G Up fwd | ATCCCCGGGTACCGAGCTCGGCCGCTCCTATGAACTTTTTG |
| ANS-C461G Up rev | ATTATCCATATCGCCATTGC |
| ANS-C461G Down fwd | GCAATGGCGATATGGATAATGGGATTGTTATTCGTTCGGCG |
| ANS-C461G Down rev | TTGTAAAACGACGGCCAGTGAGTAAGGATCATGTTGTCG |
| ANS-C461G check fwd | GCCGTACATGTTCTATATCCGTGGC |
| ANS-C461G check rev | GATCAGGCTCAACATCAGTGGC |

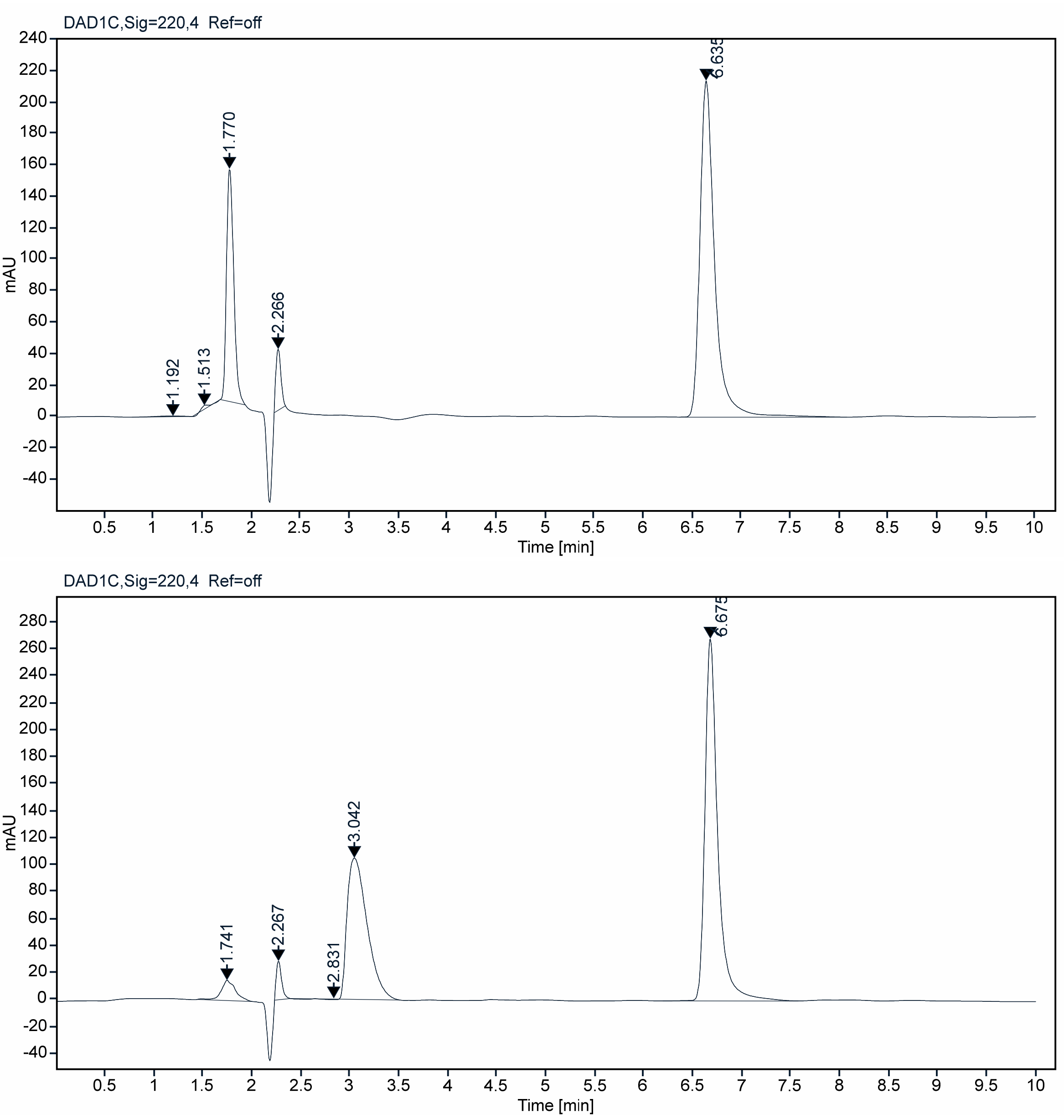


**Fig. S1A: Chromatograms of supernatant samples of *C. glutamicum* containing ANT and a by-product, presumably glycosyl-anthranilate.** The strain *C. glutamicum* ANT1 pMKEx2-$aroF^{*}_{EcCg}$-$tkt_{Cg}$ was cultivated in baffled flasks in defined CGXII medium with 4 % (w/v) glucose as carbon source and 20 µM IPTG at 30 °C and 130 RPM and samples taken during the cultivation were analyzed by HPLC**.** Top: Chromatogram at time $t$ = 0 h. The ANT peak has a retention time of 6.6 min. Bottom: Chromatogram at $t$ = 72 h. Another peak was observed at the end of the cultivation with a retention time of 2.9 min. bottom:

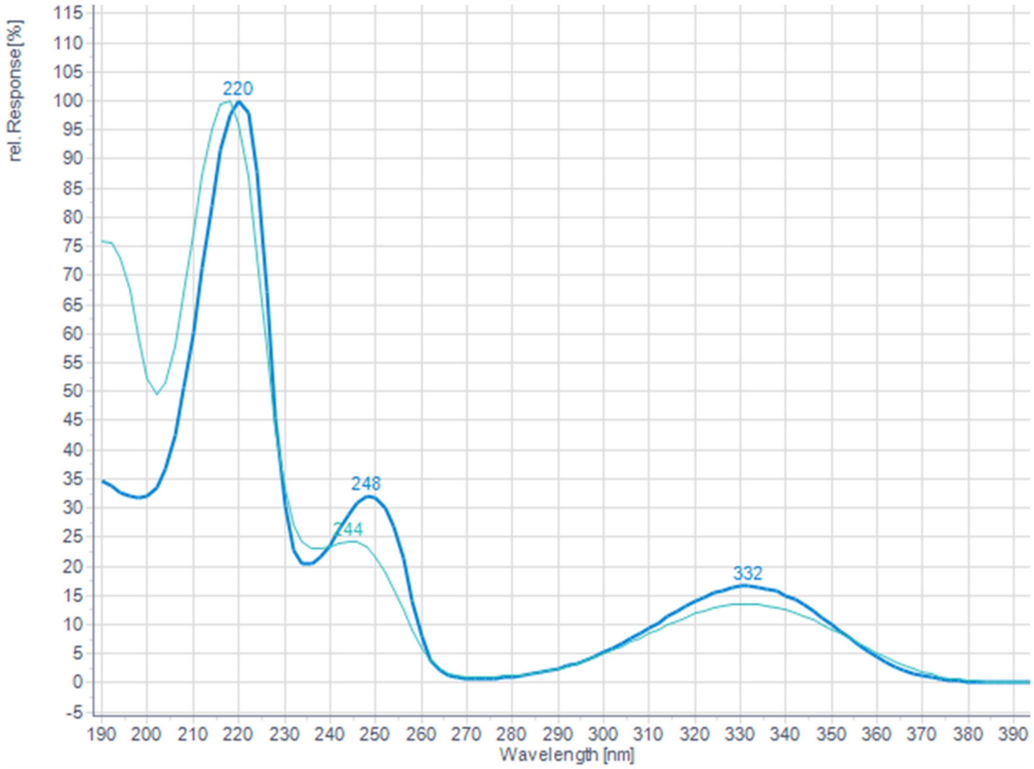


**Fig. S1B: Overlay of the UV absorption spectra of anthranilate and a by-product, presumably glycosyl-anthranilate.** Comparison of the UV absorption spectrum of the peak at 2.9 min (light blue) with the ANT peak at 6.6 min. Since the UV-absorption properties are almost identical, it was concluded that the compound with a retention time of 2.9 min is an ANT- derivative. Since this peak did not occur when ANT was dissolved in a defined CGXII medium without glucose, it was assumed to represent glycosyl-anthranilate.

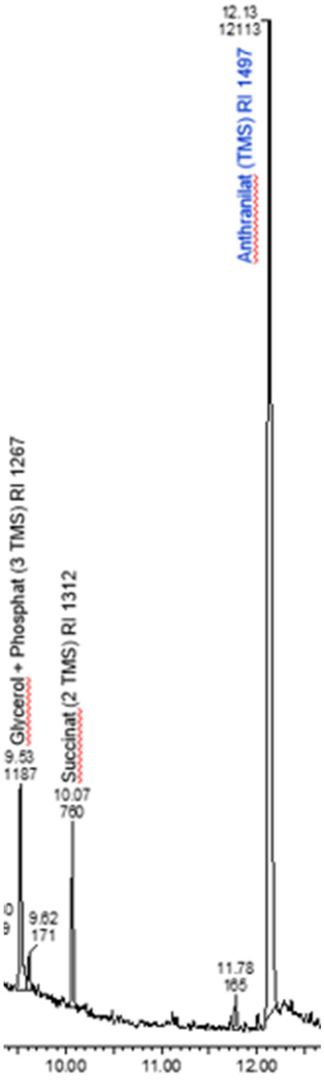


**Fig. S2: GC-TOF-MS analysis of culture supernatant of ANT-producing strain *C. glutamicum* ANT1.** Glycerol and Shikimate were identified as by-products.

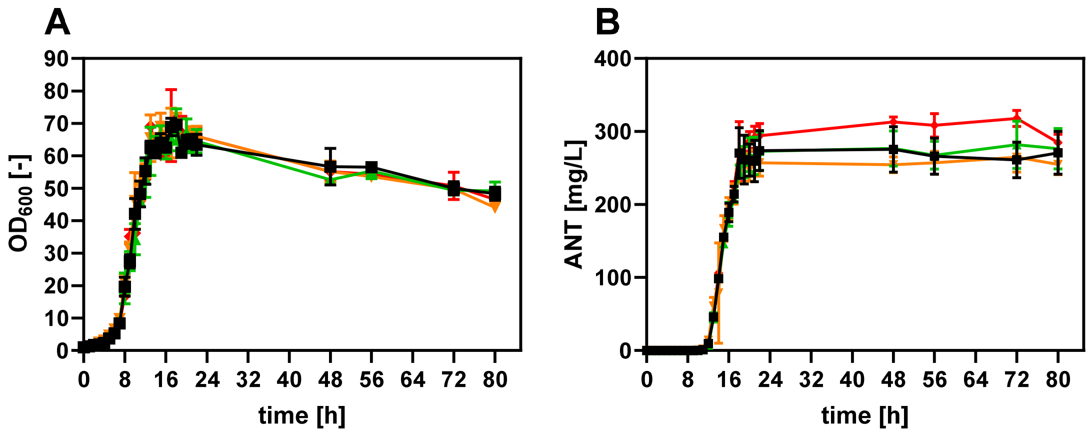


**Fig. S3: Impact of abolished competing pathways on ANT production with *C. glutamicum*.** (**A**) Growth ($OD_{600}$) of the strains *C. glutamicum* ANT1 (squares), *C. glutamicum* ANT1 Δ*qsuD* (triangles) *C. glutamicum* ANT1 Δ*nagD* (reversed triangles), *C. glutamicum* ANT1 Δ*qsuD* Δ*nagD* (ANT2, diamonds) throughout 72 h. (**B**) Determination of the ANT titer in the culture supernatants by HPLC. All strains express pMKEx2-*aroF**$_{EcCg}$-*tkt*$_{Cg}$ for ANT production from glucose. The depicted data represent mean values and standard deviation of biological triplicates.

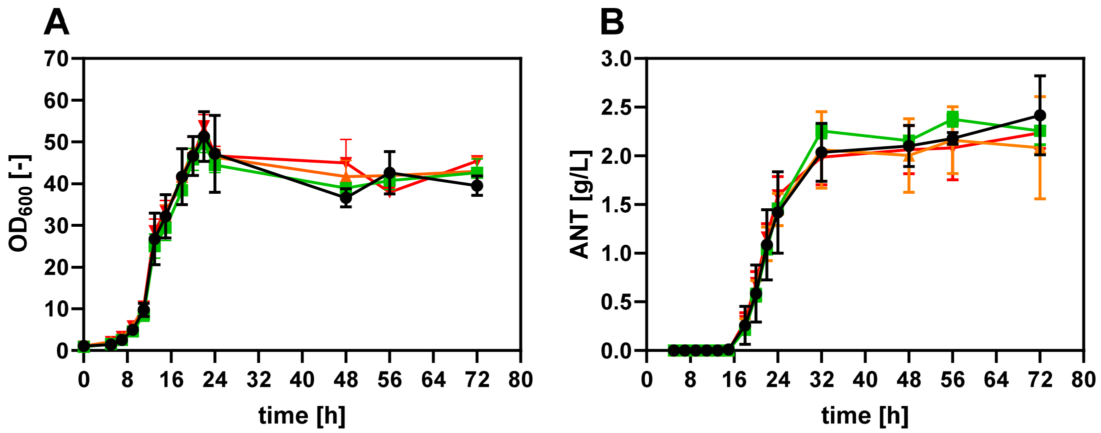


**Fig. S4: Utilization of glucose and xylose for the production of ANT with *C. glutamicum*.** (**A**) Growth ($OD_{600}$) of the strains *C. glutamicum* ANT1 Δ*qsuD* (squares) *C. glutamicum* ANT1 Δ*nagD* (triangles), *C. glutamicum* ANT1 Δ*qsuD* Δ*nagD* (ANT2, reversed triangles) throughout 72 h. (**B**) Determination of the ANT titer in the culture supernatants by HPLC. All strains express pMKEx2-*aroF*$^*_{EcCg}$-*tkt*$_{Cg}$ for ANT production and pEKEx3-*xylA*$_{Xc}$-*xylB*$_{Cg}$ for xylose utilization via the isomerase pathway. The depicted data represent mean values and standard deviation of biological triplicates.

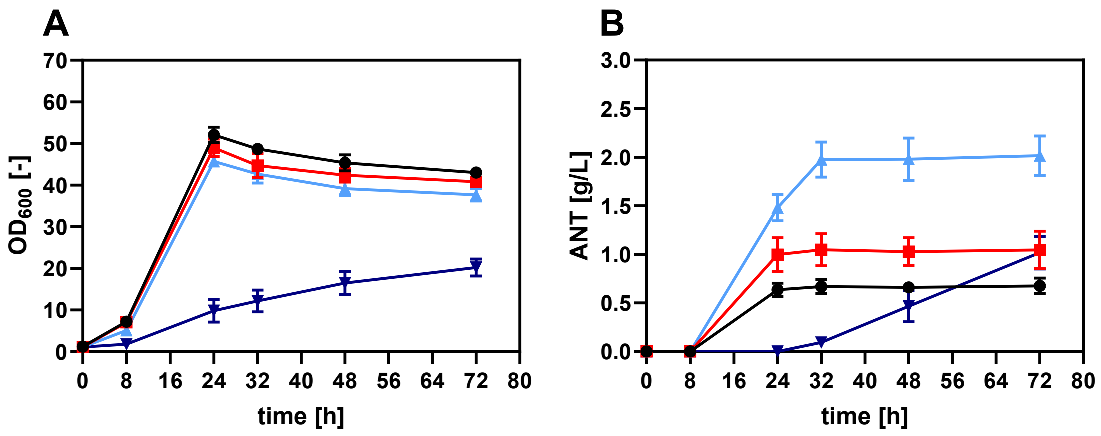


**Fig. S5**: **Impact of varying glucose/xylose ratios on the ANT titer of producing *C. glutamicum* cells.** (**A**) Growth ($OD_{600}$) of the strain *C. glutamicum* ANT2 pMKEx2-*aroF**$_{EcCg}$-*tkt*$_{Cg}$ pEKEx3-*xylA*$_{Xc}$-*xylB*$_{Cg}$ throughout 72 h. (**B**) Determination of the ANT titer in the culture supernatants by HPLC. Strains were cultivated in defined CGXII medium supplemented with 3 % glucose and 1 % xylose (circles), 2 % glucose and 2 % xylose (squares), 1 % glucose and 3 % xylose (triangles), and 4 % xylose (reversed triangles). The depicted data represent mean values and standard deviation of biological triplicates.

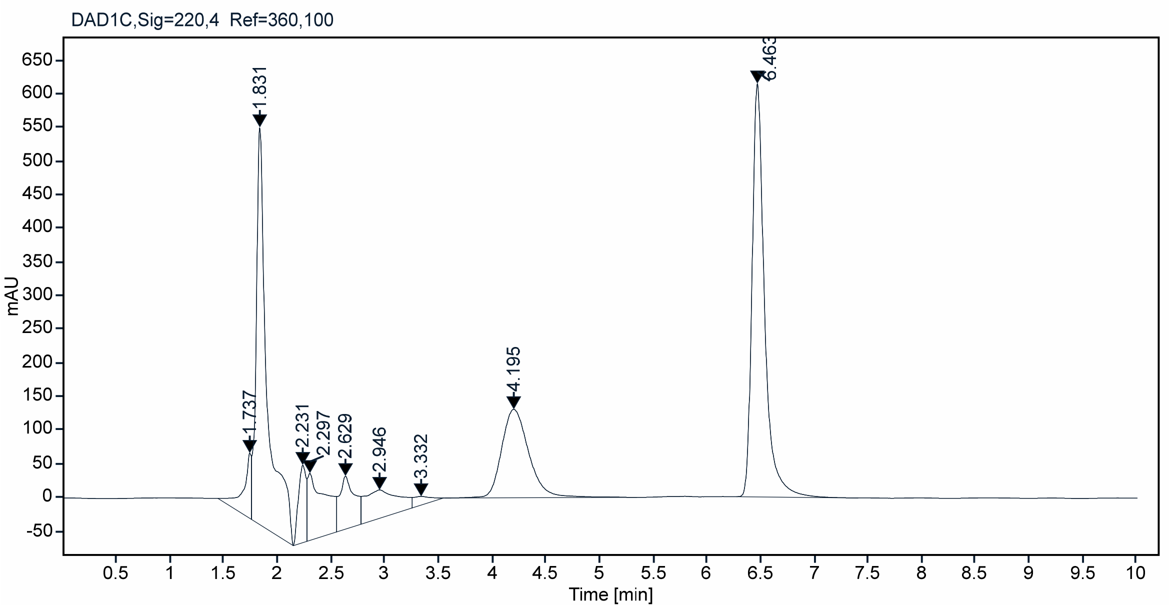


**Fig. S6: Chromatogram of *C. glutamicum* supernatant samples containing ANT and a by-product, presumably xylosyl-anthranilate.** *C. glutamicum* ANT1 pMKEx2-*aroF**$_{EcCg}$-*tkt*$_{Cg}$ pEKEx3-*xylA*$_{Xc}$-*xylB*$_{Cg}$ was cultivated in baffled flasks in defined CGXII medium with 1 % (w/v) glucose and 3 % (w/v) xylose as carbon source and 20 µM IPTG at 30 °C and 130 RPM. Samples of the supernatant were taken during the cultivation and analyzed by HPLC. The ANT peak has a retention time of 6.6 min, while the unidentified peak has a retention time of 4.3 min. Since this additional byproduct was detectable when glucose was used as the sole carbon source, it is therefore very likely that this peak represents xylosyl-anthranilate.

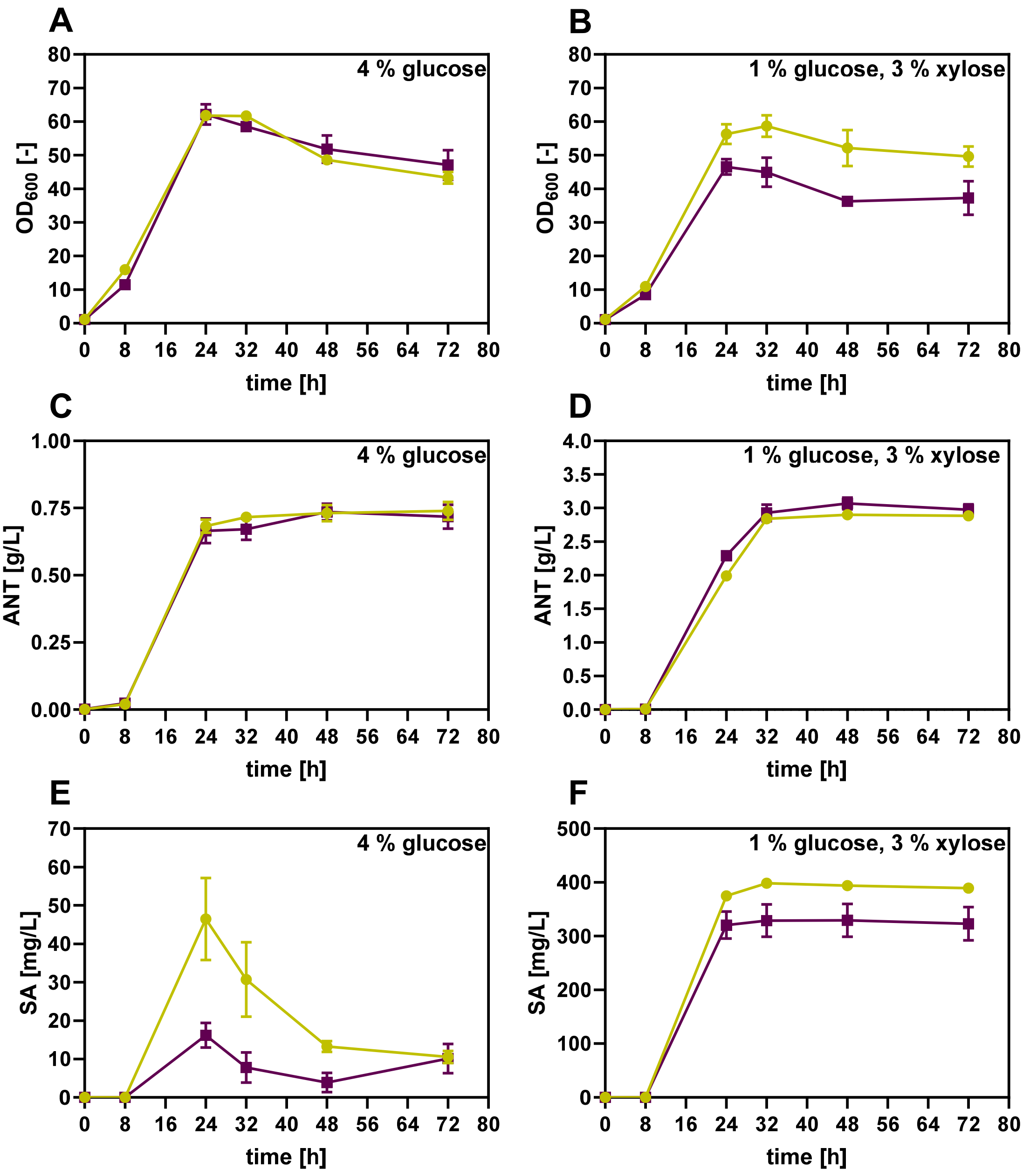


**Fig. S7: Effect of the (GTG→ATG) start codon replacement in *aroK* (shikimate kinase) on ANT and SA titer in different *C. glutamicum* strains.** (**A, B**) Growth ($OD_{600}$) on glucose or glucose/xylose mixtures of the strains *C. glutamicum* ANT3 (control, circles) and the *C. glutamicum* ANT4 variant with start codon replacement in *aroK* (GTG→ATG, squares) for 72 h. (**C, D**) Determination of the ANT titer in the culture supernatants. (**E, F**) Determination of the shikimate titers by HPLC All strains express pMKEx2-*aroF*$^*_{EcCg}$-*tkt*$_{Cg}$ for ANT production and pEKEx3-*xylA*$_{Xc}$-*xylB*$_{Cg}$ for xylose utilization via the isomerase pathway. The depicted data represent mean values and standard deviation of biological triplicates.

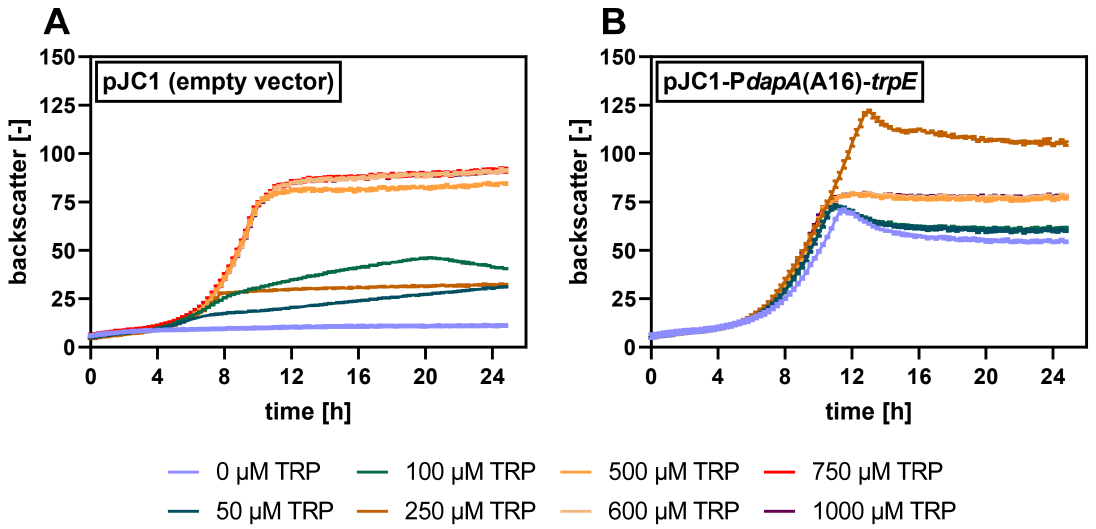


**Fig. S8: Episomal expression of *trpE* to restore growth of a *trpE*-deficient *C. glutamicum* strain in CGXII minimal medium.** (**A**) Growth of the *trpE*-deficient strain *C. glutamicum* ANT5 Δ*trpE* pJC1 harboring the empty pJC1 vector (control) in defined CGXII medium with 2 % (w/v) glucose supplemented with 0-1,000 µM TRP using a BioLector microbioreactor. (**B**) Episomal expression of *trpE* utilizing plasmid pJC1-P*dapA*(A16)-*trpE* restored growth of the *trpE*-deficient strain *C. glutamicum* ANT5 Δ*trpE* in defined CGXII medium without TRP supplementation. Biomass formation was followed by measuring the backscattered light intensity (gain 10) at a wavelength of 620 nm. The depicted data represent the mean values of biological triplicates.

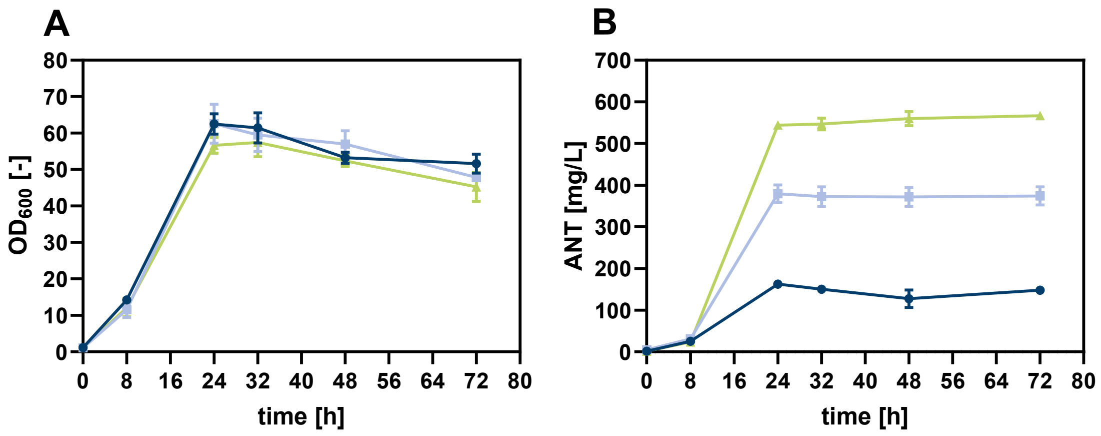


**Fig. S9**: **Genomic integration of *aroF*$^{*}_{EcCg}$ encoding DAHP synthase from *E. coli* to enable plasmid-free production of ANT.** (**A**) Growth ($OD_{600}$) of the strains *C. glutamicum* ANT4 (control, circles), *C. glutamicum* ANT4 pMKEx2-*aroF*$^{*}_{EcCg}$-*tkt*$_{Cg}$ (triangles) and *C. glutamicum* ANT4 with chromosomally integrated *aroF*$^{*}_{EcCg}$ whose expression was under the control of the *dapA* promotor variant A16 (squares) throughout 72 h. (**B**) Determination of the ANT titer in the culture supernatants. The depicted data represent mean values and standard deviation of biological triplicates.

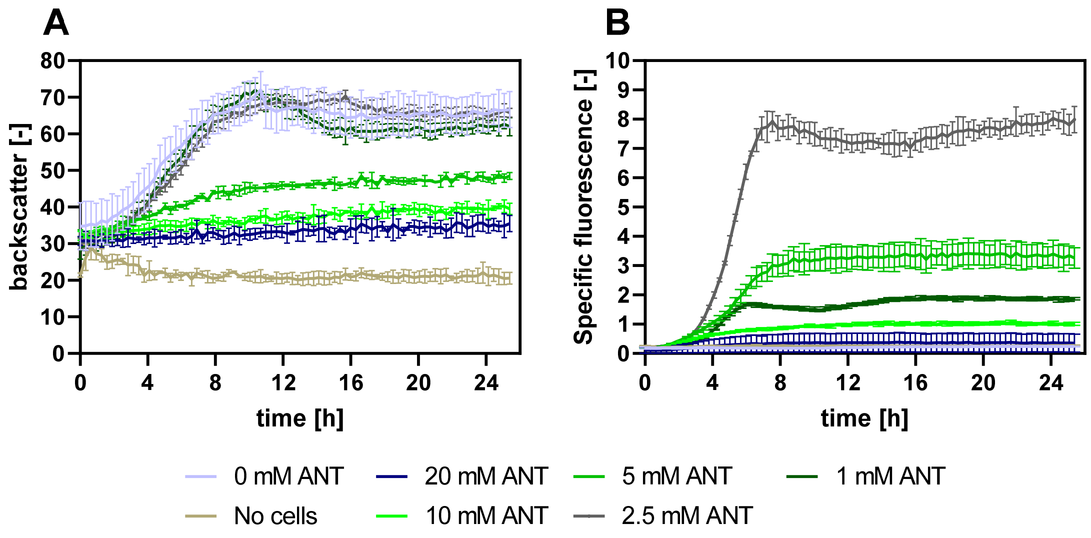


**Fig. S10: Induction of the pSen6MSA biosensor by anthranilate.** *E. coli* DH10B Δ*hcaREFCBD* harboring the 6MSA biosensor pSen6MSA was cultivated in YNB medium + 0.51 % glycerol + 2 mM Leu supplemented with 0-20 mM ANT dissolved in water using a BioLector microbioreactor. Cultures without ANT supplementation and cultivations without cells served as controls. (**A**) Biomass formation was followed by measuring the backscattered light intensity (gain 20) at a wavelength of 620 nm. (**B**) EYFP fluorescence was determined as the emission of fluorescence at 532 nm (gain 100) after excitation at 510 nm. The depicted data represent mean values and standard deviation of biological triplicates.

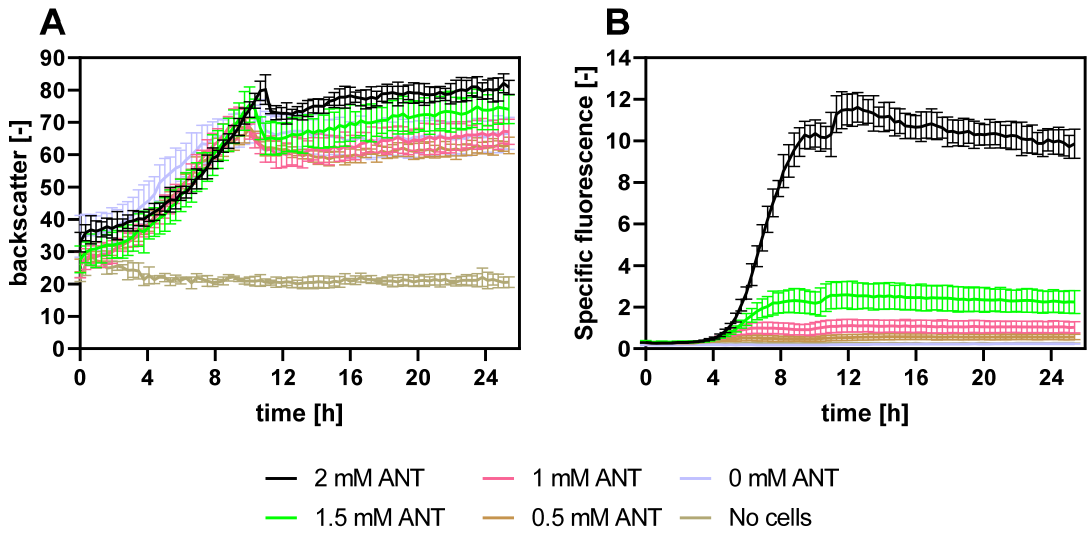


**Fig. S11: Induction of the pSen6MSA biosensor by anthranilate produced with *C. glutamicum***. *E. coli* DH10B Δ*hcaREFCBD* harboring the 6MSA biosensor pSen6MSA was cultivated in YNB medium + 0.51 % glycerol + 2 mM Leu supplemented with 0-2 mM ANT in the culture supernatant of *C. glutamicum* using a BioLector microbioreactor. Cultures without ANT supplementation and cultivations without cells served as controls. (**A**) Biomass formation was followed by measuring the backscattered light intensity (gain 20) at a wavelength of 620 nm. (**B**) EYFP fluorescence was determined as the emission of fluorescence at 532 nm (gain 100) after excitation at 510 nm. The depicted data represent mean values and standard deviation of biological triplicates.

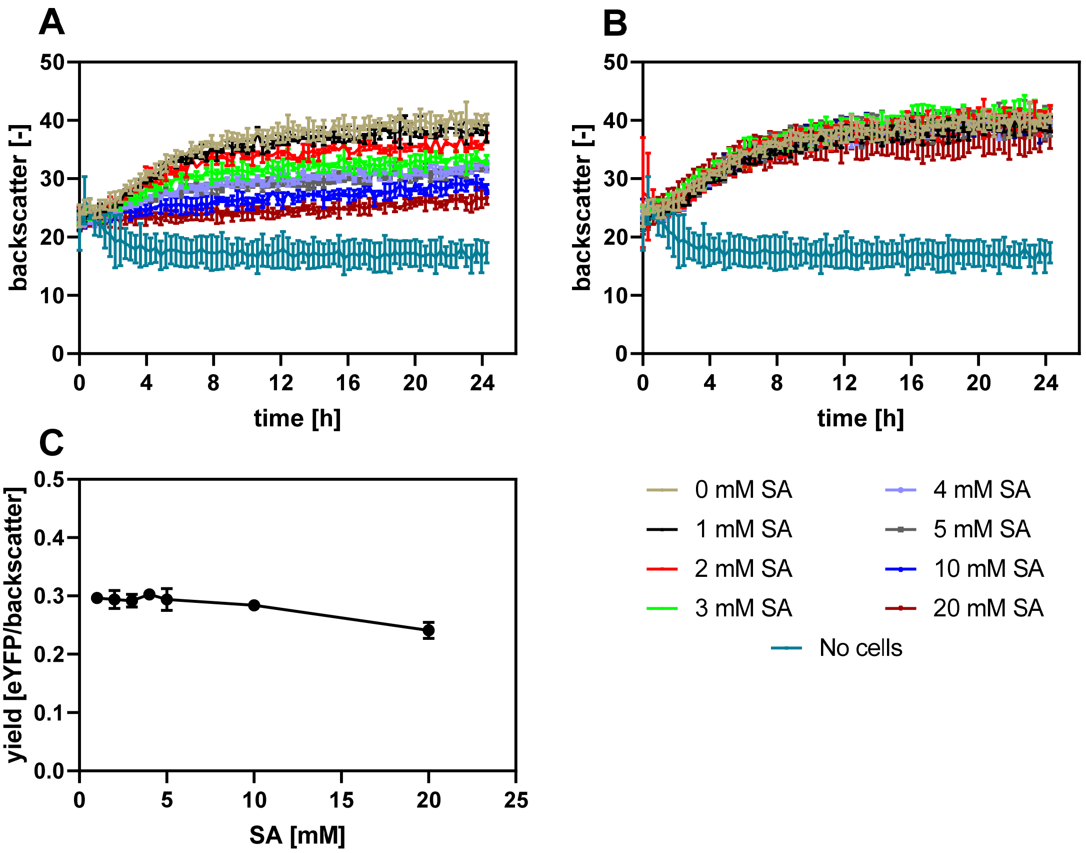


**Fig. S12: Testing shikimate as an inducer of the pSen6MSA biosensor.** *E. coli* DH10B Δ*hcaREFCBD* harboring the 6MSA biosensor pSen6MSA was cultivated in YNB medium + 0.51 % glycerol + 2 mM Leu supplemented with 0-20 mM shikimate (SA) using a BioLector microbioreactor. As controls, 0 mM SA and also no cells were cultivated. (**A**) Biomass formation was followed by measuring the backscattered light intensity (gain 20) at a wavelength of 620 nm. (**B**) EYFP fluorescence was determined as the emission of fluorescence at 532 nm (gain 100) after excitation at 510 nm. (**C**) The yield was calculated as the ratio of EYFP fluorescence and backscatter. To determine the operational range, the yield was determined for the time point t = 24 h. The depicted data represent mean values and standard deviation of biological triplicates.

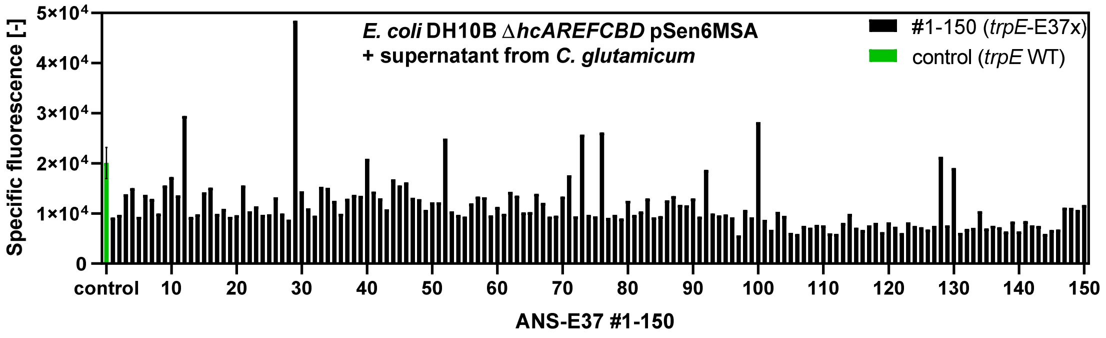


**Fig. S13: Characterization of a *C. glutamicum* strain library with randomly mutagenized nucleotide triplet encoding ANS-E37.**

**Tab. S3:** ANT titer determined for the control strains with wild-type ANS and of three clones of each of the strain library with randomly mutagenized nucleotide triplet encoding ANS-E37 with higher, equal, or lower specific fluorescence compared to the control.

| Control strains | | Higher fluorescence | | Equal fluorescence | | Lower fluorescence | |
|---|---|---|---|---|---|---|---|
| Clone | ANT [g/L] | Clone | ANT [g/] | Clone | ANT [g/L] | Clone | ANT [g/L] |
| Control 1 | 2.6 | 12 | 2.5 | 40 | 1.7 | 1 | 0.1 |
| Control 2 | 2.5 | 29 | 2.4 | 71 | 1.3 | 2 | 0.1 |
| Control 3 | 2.5 | 100 | 2.6 | 92 | 1.8 | 3 | 1 |

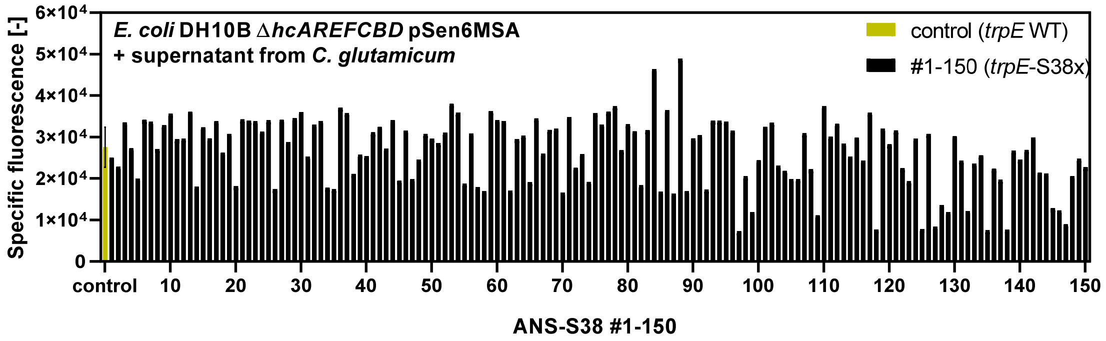


**Fig. S14: Characterization of a *C. glutamicum* strain library with randomly mutagenized nucleotide triplet encoding ANS-S38**

**Tab. S4:** ANT titer determined for the control strains with wild-type ANS and of three clones each of the strain library with randomly mutagenized nucleotide triplet encoding ANS-S38 with higher, equal, or lower specific fluorescence compared to the control.

| Control strains | | Higher fluorescence | | Equal fluorescence | | Lower fluorescence | |
|---|---|---|---|---|---|---|---|
| Clone | ANT [g/L] | Clone | ANT [g/] | Clone | ANT [g/L] | Clone | ANT [g/L] |
| Control 1 | 2.1 | 84 | 2.7 | 4 | 2 | 5 | 0.9 |
| Control 2 | 2.2 | 88 | 2.6 | 8 | 1.9 | 14 | 0 |
| Control 3 | 2.3 | 110 | 2.4 | 18 | 1.8 | 34 | 0.3 |

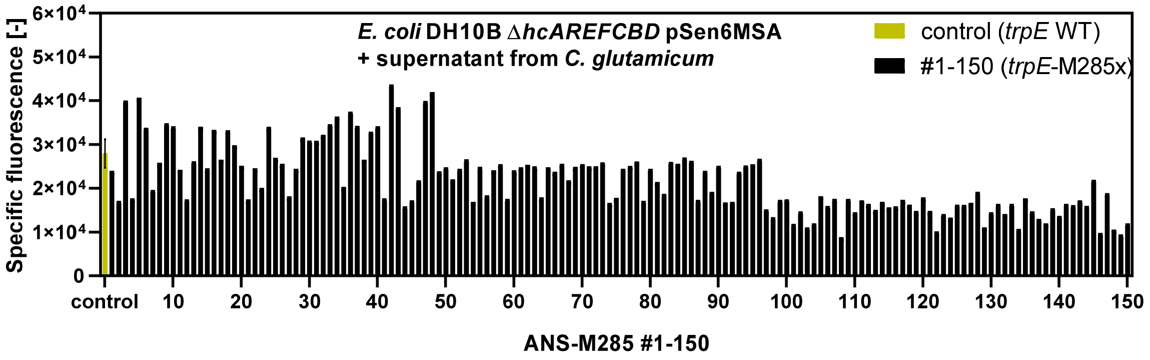


**Fig. S15 Characterization of a *C. glutamicum* strain library with randomly mutagenized nucleotide triplet encoding ANS-M285.**

**Tab. S5:** ANT titer determined for the control strains with wild-type ANS and of three clones each of the strain library with randomly mutagenized nucleotide triplet encoding ANS-M285 with higher, equal, or lower specific fluorescence compared to the control.

| Control strains | | Higher fluorescence | | Equal fluorescence | | Lower fluorescence | |
|---|---|---|---|---|---|---|---|
| Clone | ANT [g/L] | Clone | ANT [g/] | Clone | ANT [g/L] | Clone | ANT [g/L] |
| Control 1 | 2.2 | 3 | 2.1 | 19 | 1.6 | 2 | 0.1 |
| Control 2 | 2.3 | 5 | 2.2 | 96 | 1.8 | 129 | 0 |
| Control 3 | 2.2 | 42 | 1.6 | 40 | 1.6 | 149 | 0.1 |

**Tab. S6:** ANT titer determined for the control strains with wild-type ANS and of three clones each of the strain library with randomly mutagenized nucleotide triplet encoding ANS-C461 with higher, equal, or lower specific fluorescence compared to the control.

| Control strains | | Higher fluorescence | | Equal fluorescence | | Lower fluorescence | |
|---|---|---|---|---|---|---|---|
| Clone | ANT [g/L] | Clone | ANT [g/] | Clone | ANT [g/L] | Clone | ANT [g/L] |
| Control 1 | 1.9 | 29 | 1.9 | 11 | 1.8 | 24 | 0.1 |
| Control 2 | 1.6 | 44 | 2 | 19 | 2 | 87 | 0.1 |
| Control 3 | 1.8 | 135 | 2 | 97 | 1.9 | 131 | 0.1 |

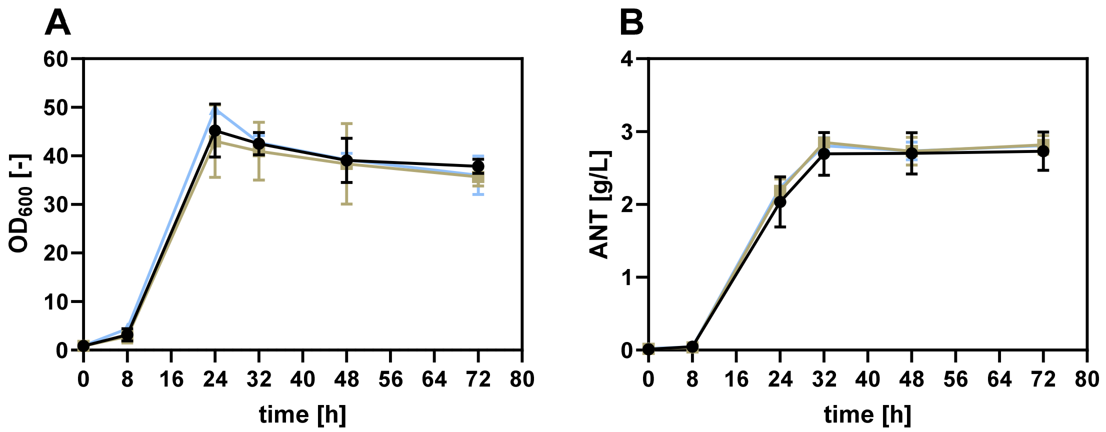


**Fig. S16: Impact of the amino acid substitution S38A and S38G in anthranilate synthase component I on ANT production with *C. glutamicum* using glucose and xylose as carbon source.** (**A**) Growth ($OD_{600}$) of the strains *C. glutamicum* ANT5 (control harboring ANS-S38R, circles), *C. glutamicum* ANT5 ANS-S38A (squares), *C. glutamicum* ANT5 ANS-S38G (triangles) harboring pMKEx2-*aroF*$_{EcCg}$-tkt$_{Cg}$* and pEKEx3-*xylA$_{Xc}$-xylB$_{Cg}$* throughout 72 h. (**B**) Determination of the ANT titer in the culture supernatants. The depicted data represent mean values and standard deviation of biological triplicates.

**Tab. S7**. KPIs of *C. glutamicum* ANT6 obtained from lab-scale batch cultivations. Maximum titers, yields and specific rates were derived via process modeling.

| Strain | *C. glutamicum* ANT6 | |
|---|---|---|
| | R1 | R2 |
| $OD_{600}$ [-] | 42.05 | 39.78 |
| CDW [g $L^{-1}$] | 10.66 | 10.09 |
| Anthranilate titer [g $L^{-1}$] | 2.47 | 2.43 |
| Growth rate from D-glucose [$h^{-1}$] | 0.38 | 0.37 |
| Growth rate from D-xylose [$h^{-1}$] | 0.08 | 0.08 |
| D-glucose uptake rate [$mmol_{Glc}$ $g_{CDW}$ $h^{-1}$] | 3.93 | 3.84 |
| D-xylose uptake rate [$mmol_{Xyl}$ $g_{CDW}$ $h^{-1}$] | 3.34 | 3.21 |
| Anthranilate production rate from D-glucose [$mmol_{Ant}$ $g_{CDW}$ $h^{-1}$] | 0.00 | 0.00 |
| Anthranilate production rate from D-xylose [$mmol_{Ant}$ $g_{CDW}$ $h^{-1}$] | 0.14 | 0.14 |
| Anthranilate yield [g $g^{-1}$][a] | 0.06 | 0.06 |

[a]Product yield per total amount of initial D-glucose and D-xylose
